# In-depth Characterization of S-Glutathionylation in Ventricular Myosin Light Chain 1 Across Species by Top-Down Proteomics

**DOI:** 10.1101/2024.12.11.628048

**Authors:** Emily A. Chapman, Holden T. Rogers, Zhan Gao, Hsin-Ju Chan, Francisco J. Alvarado, Ying Ge

## Abstract

S-glutathionylation (SSG) is increasingly recognized as a critical signaling mechanism in the heart, yet SSG modifications in cardiac sarcomeric proteins remain understudied. Here we identified SSG of the ventricular isoform of myosin light chain 1 (MLC-1v) in human, swine, and mouse cardiac tissues using top-down mass spectrometry (MS)-based proteomics. Our results enabled the accurate identification, quantification, and site-specific localization of SSG in MLC-1v across different species. Notably, the endogenous SSG of MLC-1v was observed in human and swine cardiac tissues but not in mice. Treating non-reduced cardiac tissue lysates with GSSG elevated MLC-1v SSG levels across all three species.

## Introduction

The balance between reactive oxygen species (ROS) and antioxidants is essential for maintaining redox homeostasis and regulating physiological function.^1^ Disruption of this balance, often through increased ROS exposure, leads to the reversible oxidation of reactive cysteine (Cys) thiols through various post-translational modifications (PTMs), including S-glutathionylation (SSG).^2^ SSG involves the conjugation of glutathione to a reactive protein Cys thiol, resulting in a 305.07 Da mass increase in the target protein.^3, 4^ This modification can protect protein Cys thiols from irreversible oxidation and is increasingly recognized as a critical signaling mechanism in the heart, regulating key cellular processes such as oxidative phosphorylation, cellular metabolism, and protein synthesis.^5, 6^ Cardiac sarcomeric proteins, most notably cardiac myosin-binding protein C (cMyPB-C), have been previously recognized as substrates of SSG.^5, 7, 8^ However, SSG modifications remain understudied in endogenous tissues due to their low abundance and labile nature.^5, 9, 10^ Additionally, it is unclear whether SSG is conserved across different species.

Top-down mass spectrometry (MS)-based proteomics analyzes intact proteins enabling the unambiguous identification and characterization of proteoforms - protein products from a single gene, including PTMs, alternative splice variants, and genetic mutations.^11, 12^ In contrast to traditional bottom-up MS-based proteomics, which involves proteolytic digestion of proteins into small peptides, top-down proteomics allows for the precise identification and quantification of intact proteins and their associated proteoforms directly from complex biological samples without digestion.^13, 14^ Additionally, top-down proteomics enables the fragmentation of intact proteoforms by tandem MS (MS/MS) for sequence characterization and site-localization of PTMs.^11^ As such, top-down proteomics provides an unbiased and detailed perspective on proteoform diversity and complexity.

Herein, we developed a top-down mass spectrometry (MS)-based proteomics platform to identify, quantify, and localize SSG modifications in sarcomeric proteins from human, swine, and mouse cardiac tissues. Our analysis unequivocally identified the ventricular isoform of myosin light chain 1 (MLC-1v) in human, swine, and mouse cardiac tissues as a substrate for protein SSG. Cardiac MLC-1v, also known as the essential light chain, is a key component of the hexameric myosin II complex and plays a crucial role in regulating cross-bridge cycling and cardiac contraction.^15, 16^ Recently we have demonstrated that MLC-1v, rather than MLC-2v, is ventricle-specific in adult human hearts.^17^ This study represents the first report of SSG in MLC-1v. Our results enabled the quantification and site-specific localization of SSG to Cys residues in MLC-1v across multiple species, demonstrating the power of top-down MS-based proteomics in uncovering novel PTMs within complex biological matrices.

## Materials and methods

### Detailed materials and methods are outlined in the Supporting Information

#### Chemicals and reagents

All reagents were purchased from Sigma-Aldrich unless otherwise noted. Optima LC/MS grade acetonitrile and Titan3, 17mm PES membrane syringe filters were purchased from Thermo Fisher Scientific. Amicon, 0.5 mL cellulose ultra-centrifugal filters with a molecular weight cut-off (MWCO) of 10 kDa and HPLC grade LiChrosolv® Ethanol were purchased from MilliporeSigma.

#### Human cardiac tissue collection

Left ventricular (LV) myocardium from healthy donor hearts with no history of heart disease were obtained from the University of Wisconsin (UW)-Madison Organ and Tissue Donation-Surgical Recovery and Preservation Services. The procedures for the collection of human donor heart tissues were approved by the UW-Madison Institutional Review Board (IRB). A summary of the clinical characteristics of human hearts used in this study can be found in **Table S1**.

#### Swine and mouse cardiac tissue collection

Swine heart tissue was obtained from adult Yorkshire domestic swine. Mouse heart tissue was obtained from C57Bl/6J wild-types, with an equal ratio of male to female mice. The hearts were excised from the organism, and the LV was isolated and immediately snap-frozen in liquid nitrogen and stored at -80 ºC. All experiments involving animals were conducted in accordance with the NIH Guide for the Care and Use of Laboratory Animals and using protocols approved by the University of Wisconsin Institutional Animal Care and Use Committee.

#### Non-reducing protein extraction from human, swine, and mouse cardiac tissues

Proteins were extracted from cardiac left ventricular tissue as previously reported.^18, 19^ No reducing agents were added to the extraction buffers to preserve protein SSG. Following the extraction procedure, the protein-enriched cardiac tissue lysate was passed through a Titan3, 17mm PES membrane syringe filter (pre-soaked with LiCl extraction buffer) using a 5 mL Luer-lock syringe into an Eppendorf Protein Lo-Bind tube, snap-frozen in liquid nitrogen, and stored at -80 ºC until top-down proteomics analysis.

#### Sample preparation for top-down proteomics analysis

Cardiac tissue lysates were desalted using a 10 kDa MWCO filter (Amicon, 0.5 mL, cellulose, MilliporeSigma) and buffer exchanged using 0.1% formic acid in nanopure water. For direct infusion offline MS/MS experiments, cardiac tissue lysates were buffer-exchanged twice using 0.1% formic acid in 10:10:80 IPA:ACN:water using Bio-Spin columns with Bio-Gel P-30.

#### Online top-down LC-MS/MS data acquisition

LC-MS/MS analysis was carried out using either an Acquity ultra-high performance LC M-Class system (Waters) coupled to a maXis II quadrupole time-of-flight mass spectrometer (Bruker Daltonics) or a NanoAcquity ultra-high performance LC system (Waters) coupled to an Impact II quadrupole time-of-flight mass spectrometer (Bruker Daltonics).^13, 14, 17^ 500 ng of total protein was injected onto a home-packed PLRP column (PLRP-S) (Agilent Technologies), 10 μm particle size, 250 or 500 μm inner diameter, 1,000 Å pore size using an organic gradient of 5% to 95% mobile phase B at a flow rate of 8-12 μL/min and temperature of 60 °C. Proteoforms of interest were first isolated in the gas phase and fragmented by either collisionally activated dissociation (CAD) or electron transfer dissociation (ETD).

#### Offline top-down MS/MS data acquisition

Samples were either directly infused^19^, or individual protein fractions were collected following reversed-phase LC separation for offline MS/MS analysis. Protein fractions were analyzed by nano-ESI via direct infusion using a TriVersa Nanomate system (Advion BioSciences) coupled to a 12-T solariX XR FTICR mass spectrometer (Bruker Daltonics).^20-22^ Proteoforms of interest were first isolated in the gas phase and fragmented by either collisionally activated dissociation (CAD) or electron capture dissociation (ECD).

## Data analysis

All MS data were processed and analyzed using Compass DataAnalysis v. 4.3 (Bruker Daltonics) software. MS/MS data were analyzed using MASH Native v. 1.1 software^23^ for sequence mapping and proteoform identification.

## Statistical Analysis

Paired Student’s t-tests were used to determine the level of statistical significance for the quantification of SSG modifications. All *p*-values at *p* < 0.05 were considered significant. Error bars indicated in the figures represent the mean ± standard error of the mean (SEM).

## Results & Discussion

A label-free top-down MS-based proteomics platform was used to investigate SSG modifications in sarcomeric proteins extracted directly from human (*n* = 12), swine (*n* = 6), and mouse (*n* = 8) left ventricular cardiac tissues **(Figure 1a, Figure S1)**. This platform involved tissue homogenization, extraction of proteins using a buffer without reducing agents to preserve protein SSG, and separation of intact proteins using reverse-phase liquid chromatography (RPLC). Online and offline MS/MS were then performed to comprehensively characterize protein SSG in the sarcomere **(Figure 1a, Figure S1)**. The high reproducibility of our online LC-MS/MS method was demonstrated between triplicate injections of the same amount of total protein **(Figure S2)**, enabling quantitative analysis of SSG proteoforms.

**Figure 1.**
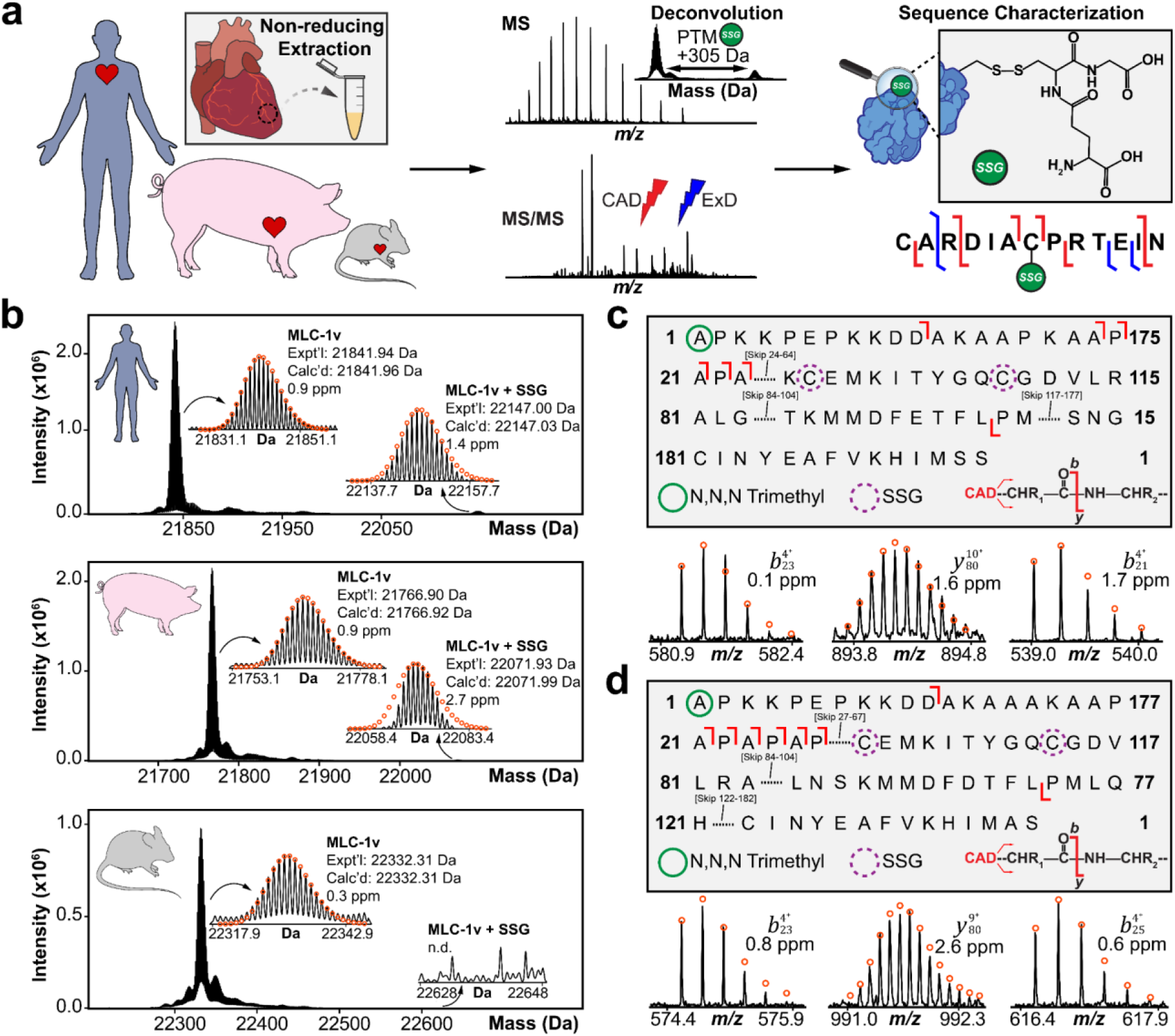
Endogenous SSG of MLC-1v is detected in human and swine cardiac tissues, but not in mouse tissues using top-down proteomics. **(a)** Top-down proteomics workflow for the identification of protein SSG (+305.07 Da) in human, swine, and mouse cardiac tissues. Intact sarcomeric proteins were extracted from cardiac tissue using a non-reducing extraction buffer. MS and MS/MS data were acquired for intact protein mass analysis, sequence characterization, and SSG localization. **(b)** Representative high-resolution deconvoluted mass spectra of MLC-1v in human, swine, and mouse cardiac tissues. Theoretical isotope distributions (red circles) are overlaid on the experimentally obtained mass spectra. The MLC-1v + SSG proteoform was isolated and fragmented in human and swine using collisionally activated dissociation (CAD). The CAD fragmentation map for **(c)** human and **(d)** swine MLC-1v shows SSG is modified on either Cys66 or Cys75 in human and Cys68 or Cys77 in swine. The green circle represents N,N,N-trimethylation, while the dashed purple circle represents possible SSG sites. Representative CAD fragment ions are shown below the fragmentation maps. All individual ion assignments are within 10 ppm of the theoretical mass and the theoretical isotopic distributions are indicated by the red circles.

### Top-Down Proteomics Reveals Endogenous SSG of MLC-1v

Our top-down proteomics data revealed low-abundant endogenous SSG on MLC-1v in human **(Figure 1b, Figure S3)** and swine **(Figure 1b, Figure S4)** tissues with high mass accuracy, indicated by a 305.07 Da mass increase from the most abundant MLC-1v proteoform **(Table S2)**. Interestingly, endogenous SSG was not detected in mouse tissues on MLC-1v **(Figure 1b, Figure S5)**. The total amount of SSG detected on MLC-1v (%SSG) was quantified by the ratio of the deconvoluted peak intensity of the SSG proteoform to the summed peak intensities of all MLC-1v proteoforms. Total SSG levels of MLC-1v in human tissues ranged from 0.2% to 1.5% across twelve biological replicates and from 0.1% to 0.4% across six biological replicates in swine tissues. No significant sex-based differences were detected in total SSG levels in our sex-balanced human cohort (6 females and 6 males) **(Figure S6)**. To ensure the reproducibility of our online LC-MS/MS method for quantification of endogenous MLC-1v SSG proteoforms, we performed three extraction replicates using human **(Figure S7)** and swine cardiac tissues **(Figure S8)**. Extraction reproducibility was demonstrated by the highly similar base peak chromatograms (BPC) and comparable %SSG values calculated for MLC-1v between extraction replicates. To localize the site of endogenous SSG in human and swine MLC-1v, we performed online LC-MS/MS with collisionally activated dissociation (CAD) to produce N- and C-terminal protein fragment ions of the isolated MLC-1v SSG proteoform **(Figure 1c-d, Figure S9-S10)**. Fragment ions confirmed N,N,N-trimethylation to Ala1 in both human and swine cardiac tissues, as previously reported.^17, 20^ However, due to the low abundance of the endogenous SSG proteoform, we could only narrow down the site of SSG to be on either Cys66 or Cys75 in human and Cys68 or Cys77 in swine, ruling out Cys181 and Cys183 as possible sites of SSG in human and swine MLC-1v, respectively. Due to the low signal-to-noise ratio of fragment ions in the online MS/MS spectra, which hindered site-localization of the SSG modification, we explored an alternative approach to further fragment and localize SSG in MLC-1v.

### Treatment with oxidized glutathione (GSSG) leads to increased SSG levels in MLC-1v

Human, swine, and mouse tissue lysates were treated with 1 mM of oxidized glutathione (GSSG) prior to top-down proteomics analysis. We observed a significant increase in the abundance of the MLC-1v SSG proteoform in human **(Figure 2a, Figure 2d, Figure S11)** and swine **(Figure 2b, Figure 2e, Figure S12)** cardiac tissues following GSSG treatment, compared to controls. Interestingly, SSG of MLC-1v was detected in mouse cardiac tissues after GSSG treatment, with total SSG levels ranging from 0.6% to 1.1% across three biological replicates **(Figure 2c, Figure 2f, Figure S13)**. To determine if GSSG treatment induced SSG of other major sarcomeric proteins detected in the LC-MS/MS analysis, we analyzed the deconvoluted mass spectra of cardiac troponin T (cTnT), cardiac troponin I (cTnI), alpha-tropomyosin (α-Tpm), troponin C (TnC), ventricular isoform of myosin light chain 2 (MLC-2v), and actin in human **(Figure S14)**, swine **(Figure S15)**, and mouse **(Figure S16)**. Evidently, we did not identify cTnT, cTnI, α-Tpm, TnC, or MLC-2v in human, swine, and mouse cardiac tissues to be modified with SSG following GSSG treatment, as we did not detect isotopically resolved proteoforms with a +305.07 Da mass shift from the most abundant proteoform with high mass accuracy. However, actin appears to be potentially modified with SSG in human, swine, and mouse cardiac tissues following incubation with 1 mM GSSG **(Figure S14-S16)**. To localize the specific sites of SSG in MLC-1v in human, swine, and mouse cardiac tissues treated with GSSG, we employed a combination of offline and online MS/MS methods using CAD, electron transfer dissociation (ETD), and electron capture dissociation (ECD). For human MLC-1v, MS/MS analysis revealed key fragment ions containing the SSG modification, including *b*_191_, *y*_168_, and *y*_171_, which closely matched the theoretical isotopic distribution with high mass accuracy **(Figure 2g, Figure S17)**. The detection of the *y*_120_ ion, which is not modified with SSG, confirmed Cys66 as the putative site of SSG in human MLC-1v. For swine MLC-1v, MS/MS analysis also revealed key fragment ions containing the SSG modification, including *b*_193_, *y*_169_, and *y*_171_ **(Figure 2h, Figure S18)**. The detection of the *y*_127_ and *y*_128_ ions, which are unmodified, confirmed Cys68 as the putative site of SSG in swine MLC-1v. Finally, MS/MS analysis of mouse MLC-1v confirmed N,N,N-trimethylation to Ala1 and ruled out Cys190 as the site of SSG modification, as C-terminal fragment ions, such as *y*_80_ and *y*_61_, were unmodified **(Figure 2i, Figure S19)**. Based on the presence of the *b*_125_ ion containing the SSG modification with high mass accuracy, Cys84 was assigned as the putative site of SSG in mouse MLC-1v. Interestingly, sequence alignment of MLC-1v revealed that Cys66 in human and Cys68 swine are conserved, whereas this Cys residue is not conserved in mouse **(Figure S20)**. Instead, Cys84 in mouse, which can be modified with SSG, is conserved with Cys77 in human and Cys75 in swine; however, these residues were found to be unlikely substrates for SSG. Therefore, these findings highlight the species-specific nature of SSG and underscore the differences in how this modification may be regulated across species.

**Figure 2.**
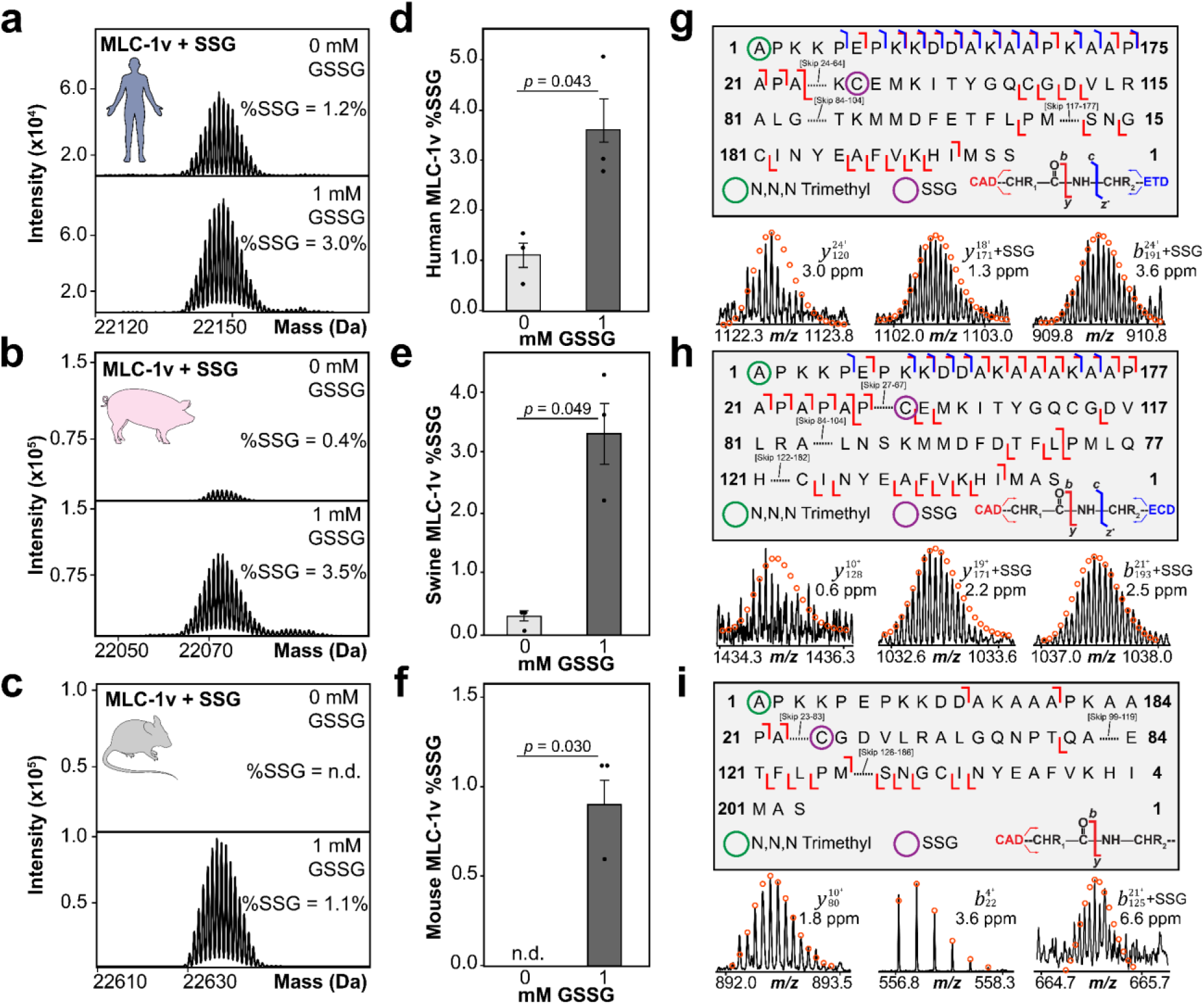
Top-down proteomics reveals treatment of non-reduced cardiac tissue lysates with GSSG increases MLC-1v SSG levels in human, swine, and mice. Representative deconvoluted mass spectra of the **(a)** human, **(b)** swine, and **(c)** mouse MLC-1v + SSG proteoform in control (0 mM GSSG) and treated (1 mM GSSG) samples. Total protein SSG (%SSG) was quantified based on the ratio of the deconvoluted peak intensity of the SSG proteoform to the summed peak intensities of all proteoforms of MLC-1v. Quantification of %SSG in **(d)** human, **(e)** swine, and **(f)** mouse MLC-1v for control (0 mM GSSG) versus treated (1 mM GSSG) samples (*n* = 3). Groups were considered significantly different by paired student’s t-test with *p* < 0.05. Error bars represent the mean ± standard error of the mean (SEM). **(g)** The collisionally activated dissociation (CAD) and electron transfer dissociation (ETD) fragmentation map for human MLC-1v (1 mM GSSG) shows SSG is localized to Cys66. The green circle represents N,N,N-trimethylation, while the purple circle represents SSG. Representative CAD fragment ions are shown below the fragmentation map. All individual ion assignments are within 10 ppm of the theoretical mass, and the theoretical isotopic distributions are indicated by the red circles. **(h)** The CAD fragmentation map for swine MLC-1v (1 mM GSSG) shows SSG is localized to Cys68. Representative CAD fragment ions are shown below the fragmentation map. **(i)** The CAD fragmentation map for mouse MLC-1v (1 mM GSSG) shows SSG is localized to Cys84. Representative CAD fragment ions are shown below the fragmentation map.

In this study, we employed top-down MS-based proteomics to identify, quantify, and localize SSG in MLC-1v from human, swine, and mouse cardiac tissues. To our knowledge, this is the first time SSG has been reported on MLC-1v. Previous studies have shown that key sarcomeric proteins, particularly cMyBP-C as well as titin, actin, and cTnI, may act as substrates for SSG, which have advanced our understanding of SSG in the sarcomere.^5, 7, 8, 24^ Titin or cMyBP-C were not detected in the current top-down 1DLC-MS data due to their large molecular weights, as in top-down proteomics, the signal-to-noise ratio (S/N) decreases with increasing protein size.^25^ Although we detected cTnI in our top-down MS data, SSG of cTnI was not observed in non-failing hearts using the non-reducing method employed in this study. A previous study has shown that SSG levels of cTnI are increased in end-stage human failing hearts compared to non-failing hearts.^5, 8^ However, other studies have not detected SSG of cTnI.^26, 27^ For example, in isolated rat control hearts, SSG was not reported on cTnI but was found to be specific to cMyBP-C.^26^ Interestingly, actin appears to be modified with SSG in human, swine, and mouse cardiac tissues upon incubation with 1 mM GSSG **(Figure S14-S16)**. Nevertheless, further effort needs to be allocated to isotopically resolve the presumable actin SSG proteoforms and subject to MS/MS to confirm this modification. In this study, we focused on the in-depth characterization of SSG in MLC-1v at the basal level and after GSSG treatment in non-failing hearts from human, swine, and mouse cardiac tissues. MLC-1v is increasingly recognized for its essential role in modulating cardiac contractility by regulating the interaction between actin and myosin within the sarcomere.^15, 16^ Recently we have demonstrated that MLC-1v, rather than MLC-2v, is exclusively expressed in the ventricles in adult human hearts, making it a potential ventricular marker.^17^ Note that SSG of MLC-1v was not detected in the previous studies^17,20^ due to the presence of reducing agents added in the extraction procedures. Here, for the first time, our non-reducing extraction procedure enabled the accurate identification, quantification, and site-specific localization of SSG in MLC-1v across different species.

## Conclusion

Overall, we identified low-abundance endogenous SSG on MLC-1v in human and swine cardiac tissues, but not in mouse tissues. Due to the low abundance of endogenous SSG on MLC-1v, samples were then treated with GSSG prior to MS analysis. Notably, SSG levels significantly increased in human and swine samples, and SSG was also detected in mouse samples following GSSG treatment. Additionally, MS/MS analysis confirmed the putative SSG sites in MLC-1v to be Cys66 in human, Cys68 in swine, and Cys84 in mouse. Overall, our results provide the first evidence that MLC-1v is a target of SSG in the sarcomere across different species, demonstrating the utility of top-down MS-based proteomics in identifying, quantifying, and localizing novel PTMs directly from cardiac tissues.

## Supporting information

Supplementary Data 1. SSG Quantification Data

Supplementary Data 2. Top-Down MSMS Results

## Acknowledgements

This work was supported by NIH R01 HL109810 (to Y.G.). Y.G. would also like to acknowledge R01 GM117058, R01 HL156855, and S10 OD018475. E.A.C. would like to acknowledge support from the NIH Chemistry-Biology Interface Training Program NIH T32GM152341. H.T.R. would like to acknowledge support from the National Heart, Lung, and Blood Institute of the NIH under Award Number T32HL007936 through the UW-Madison Cardiovascular Research Center. F.J.A. would like to acknowledge NIH R01 HL161070 and R01 HL167195.

## Data Availability

The mass spectrometry proteomics data generated in this study have been deposited to the ProteomeXchange Consortium via the PRIDE partner repository under the accession code PXD058703 and MassIVE repository under the accession code MSV000096622.

## Contributions

E.A.C., H.T.R., Z.G., H.C., and Y.G. designed research; E.A.C., H.T.R., Z.G., and R.C. performed MS analysis; E.A.C. prepared the biological samples; E.A.C. and Y.G. analyzed data; F.J.A. provided biological samples; and E.A.C., and Y.G. wrote the paper with comments from other co-authors.

## References

(1) Chung, H. S.; Wang, S.-B.; Venkatraman, V.; Murray, C. I.; Van Eyk, J. E. Cysteine Oxidative Posttranslational Modifications. Circulation Research 2013, 112 (2), 382–392. DOI: 10.1161/CIRCRESAHA.112.268680.

(2) Rashdan, N. A.; Shrestha, B.; Pattillo, C. B. S-glutathionylation, friend or foe in cardiovascular health and disease. Redox Biology 2020, 37, 101693. DOI: 10.1016/j.redox.2020.101693.

(3) Li, X.; Zhang, T.; Day, N. J.; Feng, S.; Gaffrey, M. J.; Qian, W.-J. Defining the S-Glutathionylation Proteome by Biochemical and Mass Spectrometric Approaches. In Antioxidants, 2022; Vol. 11.

(4) Kukulage, D. S. K.; Matarage Don, N. N. J.; Ahn, Y.-H. Emerging chemistry and biology in protein glutathionylation. Current Opinion in Chemical Biology 2022, 71, 102221. DOI: 10.1016/j.cbpa.2022.102221.

(5) Rosas, P. C.; Solaro, R. J. Implications of S-glutathionylation of sarcomere proteins in cardiac disorders, therapies, and diagnosis. Frontiers in Cardiovascular Medicine 2023, 9, Systematic Review.

(6) Pastore, A.; Piemonte, F. Protein Glutathionylation in Cardiovascular Diseases. In International Journal of Molecular Sciences, 2013; Vol. 14, pp 20845–20876.

(7) Patel, B. G.; Wilder, T.; Solaro, R. J. Novel control of cardiac myofilament response to calcium by S-glutathionylation at specific sites of myosin binding protein C. Frontiers in Physiology 2013, 4.

(8) Budde, H.; Hassoun, R.; Tangos, M.; Zhazykbayeva, S.; Herwig, M.; Varatnitskaya, M.; Sieme, M.; Delalat, S.; Sultana, I.; Kolijn, D.; et al. The Interplay between S-Glutathionylation and Phosphorylation of Cardiac Troponin I and Myosin Binding Protein C in End-Stage Human Failing Hearts. In Antioxidants, 2021; Vol. 10.

(9) Li, X.; Gluth, A.; Zhang, T.; Qian, W.-J. Thiol redox proteomics: Characterization of thiol-based post-translational modifications. PROTEOMICS 2023, 23 (13-14), 2200194. DOI: 10.1002/pmic.202200194.

(10) Karpov, O. A.; Stotland, A.; Raedschelders, K.; Chazarin, B.; Ai, L.; Murray, C. I.; Van Eyk, J. E. Proteomics of the heart. Physiological Reviews 2024, 104 (3), 931–982. DOI: 10.1152/physrev.00026.2023.

(11) Roberts, D. S.; Loo, J. A.; Tsybin, Y. O.; Liu, X.; Wu, S.; Chamot-Rooke, J.; Agar, J. N.; Paša-Tolić, L.; Smith, L. M.; Ge, Y. Top-down proteomics. Nature Reviews Methods Primers 2024, 4 (1), 38. DOI: 10.1038/s43586-024-00318-2.

(12) Smith, L. M.; Kelleher, N. L.; Linial, M.; Goodlett, D.; Langridge-Smith, P.; Ah Goo, Y.; Safford, G.; Bonilla *, L.; Kruppa, G.; Zubarev, R.; et al. Proteoform: a single term describing protein complexity. Nature Methods 2013, 10 (3), 186–187. DOI: 10.1038/nmeth.2369.

(13) Chapman, E. A.; Aballo, T. J.; Melby, J. A.; Zhou, T.; Price, S. J.; Rossler, K. J.; Lei, I.; Tang, P. C.; Ge, Y. Defining the Sarcomeric Proteoform Landscape in Ischemic Cardiomyopathy by Top-Down Proteomics. Journal of Proteome Research 2023, 22 (3), 931–941. DOI: 10.1021/acs.jproteome.2c00729.

(14) Tucholski, T.; Cai, W.; Gregorich, Z. R.; Bayne, E. F.; Mitchell, S. D.; Mcilwain, S. J.; De Lange, W. J.; Wrobbel, M.; Karp, H.; Hite, Z.; et al. Distinct hypertrophic cardiomyopathy genotypes result in convergent sarcomeric proteoform profiles revealed by top-down proteomics. Proceedings of the National Academy of Sciences 2020, 117 (40), 24691–24700. DOI: 10.1073/pnas.2006764117.

(15) Sitbon, Y. H.; Yadav, S.; Kazmierczak, K.; Szczesna-Cordary, D. Insights into myosin regulatory and essential light chains: a focus on their roles in cardiac and skeletal muscle function, development and disease. Journal of Muscle Research and Cell Motility 2020, 41 (4), 313–327. DOI: 10.1007/s10974-019-09517-x.

(16) Hernandez, O. M.; Jones, M.; Guzman, G.; Szczesna-Cordary, D. Myosin essential light chain in health and disease. American Journal of Physiology-Heart and Circulatory Physiology 2007, 292 (4), H1643–H1654. DOI: 10.1152/ajpheart.00931.2006.

(17) Bayne, E. F.; Rossler, K. J.; Gregorich, Z. R.; Aballo, T. J.; Roberts, D. S.; Chapman, E. A.; Guo, W.; Palecek, S. P.; Ralphe, J. C.; Kamp, T. J.; et al. Top-down proteomics of myosin light chain isoforms define chamber-specific expression in the human heart. Journal of Molecular and Cellular Cardiology 2023, 181, 89–97. DOI: 10.1016/j.yjmcc.2023.06.003.

(18) Chapman, E. A.; Roberts, D. S.; Tiambeng, T. N.; Andrews, J.; Wang, M.-D.; Reasoner, E. A.; Melby, J. A.; Li, B. H.; Kim, D.; Alpert, A. J.; et al. Structure and dynamics of endogenous cardiac troponin complex in human heart tissue captured by native nanoproteomics. Nature Communications 2023, 14 (1), 8400. DOI: 10.1038/s41467-023-43321-z.

(19) Chapman, E. A.; Li, B. H.; Krichel, B.; Chan, H.-J.; Buck, K. M.; Roberts, D. S.; Ge, Y. Native Top-Down Mass Spectrometry for Characterizing Sarcomeric Proteins Directly from Cardiac Tissue Lysate. Journal of the American Society for Mass Spectrometry 2024, 35 (4), 738–745. DOI: 10.1021/jasms.3c00430.

(20) Gregorich, Z. R.; Cai, W.; Lin, Z.; Chen, A. J.; Peng, Y.; Kohmoto, T.; Ge, Y. Distinct sequences and post-translational modifications in cardiac atrial and ventricular myosin light chains revealed by top-down mass spectrometry. Journal of Molecular and Cellular Cardiology 2017, 107, 13–21. DOI: 10.1016/j.yjmcc.2017.04.002.

(21) Tucholski, T.; Knott, S. J.; Chen, B.; Pistono, P.; Lin, Z.; Ge, Y. A Top-Down Proteomics Platform Coupling Serial Size Exclusion Chromatography and Fourier Transform Ion Cyclotron Resonance Mass Spectrometry. Analytical Chemistry 2019, 91 (6), 3835–3844. DOI: 10.1021/acs.analchem.8b04082.

(22) Peng, Y.; Gregorich, Z. R.; Valeja, S. G.; Zhang, H.; Cai, W.; Chen, Y.-C.; Guner, H.; Chen, A. J.; Schwahn, D. J.; Hacker, T. A.; et al. Top-down Proteomics Reveals Concerted Reductions in Myofilament and Z-disc Protein Phosphorylation after Acute Myocardial Infarction*. Molecular & Cellular Proteomics 2014, 13 (10), 2752–2764. DOI: 10.1074/mcp.M114.040675.

(23) Larson, E. J.; Pergande, M. R.; Moss, M. E.; Rossler, K. J.; Wenger, R. K.; Krichel, B.; Josyer, H.; Melby, J. A.; Roberts, D. S.; Pike, K.; et al. MASH Native: A Unified Solution for Native Top-Down Proteomics Data Processing. Bioinformatics 2023, btad359. DOI: 10.1093/bioinformatics/btad359.

(24) Alegre-Cebollada, J.; Kosuri, P.; Giganti, D.; Eckels, E.; Rivas-Pardo Jaime A.; Hamdani, N.; Warren Chad M.; Solaro, R. J.; Linke Wolfgang A.; Fernández Julio M. S-Glutathionylation of Cryptic Cysteines Enhances Titin Elasticity by Blocking Protein Folding. Cell 2014, 156 (6), 1235–1246. DOI: 10.1016/j.cell.2014.01.056.

(25) Cai, W.; Tucholski, T.; Chen, B.; Alpert, A. J.; McIlwain, S.; Kohmoto, T.; Jin, S.; Ge, Y. Top-Down Proteomics of Large Proteins up to 223 kDa Enabled by Serial Size Exclusion Chromatography Strategy. Analytical Chemistry 2017, 89 (10), 5467–5475. DOI: 10.1021/acs.analchem.7b00380.

(26) Chakouri, N.; Reboul, C.; Boulghobra, D.; Kleindienst, A.; Nottin, S.; Gayrard, S.; Roubille, F.; Matecki, S.; Lacampagne, A.; Cazorla, O. Stress-induced protein S-glutathionylation and phosphorylation crosstalk in cardiac sarcomeric proteins - Impact on heart function. International Journal of Cardiology 2018, 258, 207–216. DOI: 10.1016/j.ijcard.2017.12.004.

(27) Wilder, T.; Ryba, D. M.; Wieczorek, D. F.; Wolska, B. M.; Solaro, R. J. N-acetylcysteine reverses diastolic dysfunction and hypertrophy in familial hypertrophic cardiomyopathy. American Journal of Physiology-Heart and Circulatory Physiology 2015, 309 (10), H1720–H1730. DOI: 10.1152/ajpheart.00339.2015.

