## Supplementary Information for "In-depth Characterization of S-Glutathionylation in Ventricular Myosin Light Chain 1 Across Species by Top-Down Proteomics"

### Table of Contents

**Supplementary Methods**

Non-reducing protein extraction from human, swine, and mouse cardiac tissues. …......S-5

Sample preparation for top-down proteomics analysis………………………….. …......S-5

Online top-down LC-MS/MS data acquisition.. …....................................................S-6

Offline top-down MS/MS data acquisition. …...........................................................S-7

Data analysis. …...........................................................................................................S-7

**Supplementary Tables**

Supplementary Table 1. Clinical characteristics of human hearts used…....…………...S-9

Supplementary Table 2. MLC-1v proteoforms identified in top-down MS experiments from human, swine, and mouse cardiac tissues….……………………......……………..S-10

**Supplementary Figures**

Supplementary Figure 1. Top-down MS-based proteomics workflow for the identification of protein SSG in human, swine, and mouse cardiac tissues……………………………………………………………………………….S-11

Supplementary Figure 2. Highly-reproducible top-down online LC-MS/MS method……………………………………………………………………..…………..S-12

Supplementary Figure 3. Top-down proteomics identifies endogenous SSG in human MLC-1v……………………………………………………………….…………….…S-13

Supplementary Figure 4. Top-down proteomics identifies endogenous SSG in swine MLC-1v………………………………………………………………………………....S-14

Supplementary Figure 5. Endogenous SSG is not detected by top-down proteomics in mouse MLC-1v..........................................................................................................S-15

Supplementary Figure 6. Sex-based differences of SSG in human MLC-1v………......S-16

Supplementary Figure 7. Reproducible online LC-MS/MS analysis of MLC-1v from human cardiac tissue……………………………………………………….……......S-17

Supplementary Figure 8. Reproducible online LC-MS/MS analysis of MLC-1v from swine cardiac tissue………………………………………………………………….……......S-18

Supplementary Figure 9. Characterization of endogenous human MLC-1v + SSG using top-down proteomics……………………..……………………………….……….......S-19

Supplementary Figure 10. Characterization of endogenous swine MLC-1v + SSG using top-down proteomics……………………………...…………………….……….......S-20

Supplementary Figure 11. Top-down proteomics reveals treatment of non-reduced human cardiac tissue lysates with GSSG increases SSG levels in MLC-1v………………......S-21

Supplementary Figure 12. Top-down proteomics reveals treatment of non-reduced swine cardiac tissue lysates with GSSG increases SSG levels in MLC-1v………………….......S-22

Supplementary Figure 13. Top-down proteomics reveals treatment of non-reduced mouse cardiac tissue lysates with GSSG increases SSG levels in MLC-1v…...……………........S-23

Supplementary Figure 14. Overview of major cardiac proteoforms accessed by top-down proteomics when non-reduced human tissue lysate was incubated with 1 mM GSSG ..S-24

Supplementary Figure 15. Overview of major cardiac proteoforms accessed by top-down proteomics when non-reduced swine tissue lysate was incubated with 1 mM GSSG ...S-25

Supplementary Figure 16. Overview of major cardiac proteoforms accessed by top-down proteomics when non-reduced mouse tissue lysate was incubated with 1 mM GSSG ..S-26

Supplementary Figure 17. Characterization of the human MLC-1v + SSG proteoform following incubation with 1 mM GSSG using top-down proteomics……………. ......S-27

Supplementary Figure 18. Characterization of the swine MLC-1v + SSG proteoform following incubation with 1 mM GSSG using top-down proteomics……………. ......S-28

Supplementary Figure 19. Characterization of the mouse MLC-1v + SSG proteoform following incubation with 1 mM GSSG using top-down proteomics……………. ......S-29

Supplementary Figure 20. Sequence alignment of human, swine, and mouse MLC-1v………………………………………………………………………………. ......S-30

**Supplementary Methods**

**Non-reducing protein extraction from human, swine, and mouse cardiac tissues.** Proteins were extracted from cardiac tissues as previously described.[1, 2] Cardiac left ventricular tissue (50-500 mg) was homogenized on ice using a Polytron homogenizer in 10 volumes (mL/g tissues) of phosphate wash buffer (5 mM $NaH_2PO_4$, 5 mM $Na_2HPO_4$ (pH 7.0), 100 mM NaCl, 125 mM L-Methionine (pH 7.5), 1 mM PMSF and 1X Halt$^{TM}$ protease and phosphatase inhibitor cocktail. The homogenate was centrifuged at 17,000 × *g* for 3 minutes at 4 ºC, and the supernatant was discarded. The washing and homogenization step was repeated, and the supernatant was once again discarded. The protein pellet was extracted using 5 volumes (mL/g tissues) of a lithium chloride (LiCl) buffer (25 mM Tris (pH 7.5), 700 mM LiCl, 125 L-Methionine (pH 7.5), 1 mM PMSF and 1X Halt$^{TM}$ protease and phosphatase inhibitor cocktail). The resulting homogenate was incubated at 4 ºC for 10 min and then centrifuged at 17,000 × *g* for 3 minutes at 4 ºC. The supernatant was transferred to new Eppendorf Protein Lo-Bind tubes and centrifuged at 21,000 × *g* for 30 min at 4 ºC to further clarify the tissue lysates. The resulting supernatant was passed through a Titan3, 17mm PES membrane syringe filter (pre-soaked with LiCl buffer) using a 5 mL Luer-lock syringe, into a new Eppendorf Protein Lo-Bind tube, snap-frozen in liquid nitrogen, and stored at -80 ºC.

**Sample preparation for top-down proteomics analysis.** Cardiac tissue lysates were thawed out on ice for 15 minutes, desalted using a 10 kDa MWCO filter (Amicon, 0.5 mL, cellulose, MilliporeSigma), and buffer exchanged using 0.1% formic acid in nanopure water. The concentration of the desalted tissue lysates was determined by the Bradford protein assay. For direct infusion offline MS/MS experiments, non-reduced cardiac tissue lysates were buffer-exchanged twice using 0.1% formic acid in 10:10:80 IPA:ACN:water using Bio-Spin columns with Bio-Gel P-30, following the protocol detailed from Bio-Rad.

**Online top-down LC-MS/MS data acquisition.** LC-MS/MS analysis was carried out using either an Acquity ultra-high performance LC M-Class system (Waters) coupled to a maXis II quadrupole time-of-flight mass spectrometer (Bruker Daltonics) or a NanoAcquity ultra-high performance LC system (Waters) coupled to an Impact II quadrupole time-of-flight mass spectrometer (Bruker Daltonics). 500 ng of total protein was injected onto a home-packed PLRP column (PLRP-S) (Agilent Technologies), 10 μm particle size, 250 or 500 μm inner diameter, 1,000 Å pore size using an organic gradient of 5% to 95% mobile phase B (mobile phase A: 0.1% formic acid in water; mobile phase B: 0.1% formic acid in 50:50 acetonitrile:ethanol) at a flow rate of 8-12 μL/min and temperature of 60 ℃. Mass spectra were acquired at a scan rate of 0.5 Hz over 200-3000 *m/z*. For the electrospray ionization source, the end plate offset, capillary, nebulizer, dry gas, and dry temp were set at 500 V, 4500 V, 0.5 bar, 4.0 L/min, and 200 °C, respectively. For tune settings, the Funnel 1 RF, isCID energy, multipole RF (for maXis II), Funnel 2 RF (for Impact II), Hexapole RF (for Impact II), quadrupole ion energy, collision energy, collision RF, transfer time, and pre-pulse storage were set at 400 Vpp, 10 eV, 400 Vpp, 400 Vpp, 400 Vpp, 5 eV, 5 eV, 2000 Vpp, 100 µs, and 15 µs, respectively. Proteoforms of interest were first isolated in the gas phase and fragmented by either collisionally activated dissociation (CAD) or electron transfer dissociation (ETD). For targeted CAD MS/MS experiments (Impact II), the precursor ion was set with a width of at least 5 *m*/*z* and mass spectra were acquired at a scan rate of 1.0 Hz over 200 – 2000 *m/z*. Collision energies were set to values ranging from 8 to 20 eV. For targeted ETD MS/MS experiments (maXis II), 3,4 hexanedione was used as the ETD reagent. The precursor ion was set with a width of at least 5 *m/z* and mass spectra were acquired at a scan rate of 1.0 Hz over 200-3000 *m/z*. The analyte accumulation time was set to 1000 ms with a 30 ms reagent injection time, and extended reaction time varied from 10 to 100 ms.

**Offline top-down MS/MS data acquisition.** Samples were either directly infused[2], or individual protein fractions were collected following reversed-phase LC separation for offline MS/MS analysis. Protein fractions were analyzed by nano-ESI via direct infusion using a TriVersa Nanomate system (Advion BioSciences) coupled to a 12-T solariX XR FTICR mass spectrometer (Bruker Daltonics).[3-5] For the nano-ESI source using a TriVersa Nanomate, the desolvating gas pressure was maintained between 0.7 and 0.8 PSI and the voltage was set to 1.7–1.8 kV. The source dry gas flow rate was set to 3 L/min at 180 °C. For the source optics, the capillary exit, deflector plate, funnel 1, skimmer voltage, funnel RF amplitude, octopole frequency, octopole RF amplitude, collision cell RF frequency, collision cell energy, and collision cell RF amplitude were set at 240.0 V, 220.0 V, 100.0 V, 50.0 V, 300.0 Vpp, 5 MHz, 600.0 Vpp, 2.0 MHz, 4V, and 2000.0 Vpp, respectively. Mass spectra were acquired with an acquisition size of 1 to 2M in the mass range between 200 and 3000 *m*/*z*. Ions were accumulated in the collision cell for 1 to 15 s and a time-of-flight of 1.000 ms was used for their transfer to the ICR cell. For offline CAD MS/MS experiments, the collisional energy was varied between 10 and 25 V. For offline electron-capture dissociation (ECD) experiments, ions were pulsed for 20 ms with a lens bias of 1.5 V and a lens voltage of 15 V.

**Data analysis.** All MS data were processed and analyzed using Compass DataAnalysis v. 4.3 (Bruker Daltonics) software. All chromatograms were smoothened by the Gauss algorithm with a smoothing width of 2.04 s. The Sophisticated Numerical Annotation Procedure (SNAP) peak-picking algorithm (quality factor: 0.4; signal-to-noise ratio (S/N): 3.0) was applied to determine the monoisotopic mass of all detected ions. Mass spectra were deconvoluted using the Maximum Entropy algorithm within the DataAnalysis v. 4.3 software. The resolving power for Maximum Entropy deconvolution was set to 50,000k (Impact II) and 60,000k (maXis II) for proteins that were isotopically resolved. Protein S-glutathionylation (SSG) was quantified based on the ratio of the peak intensity of the SSG proteoform to the summed peak intensities of all proteoforms of the same protein using the deconvoluted mass spectrum. Online and offline MS/MS data were output from the DataAnalysis v. 4.3 software and analyzed using MASH Native v. 1.1 software[6] using the eThrash algorithm[7] for sequence mapping and proteoform identification. All fragment ions were manually validated using a 10-ppm mass tolerance.

### Supplementary Tables

**Table S1. Clinical characteristics of human hearts used.** Non-failing donor human heart tissues were obtained from the University of Wisconsin (UW) Organ and Tissue Donation, Surgical Recovery, and Preservation Services. Average and standard error of the mean (SEM) is reported for age. Number of donors (*n*) is indicated in parentheses after each reported percentage.

| Clinical Characteristic | Precentage of Sample Cohort |
|---|---|
| **Age** | 55 ± 11.5 years |
| | min: 25 years |
| | max: 65 years |
| **Male** | 50% (*n* = 6) |
| **Female** | 50% (*n* = 6) |

**Table S2. MLC-1v proteoforms identified in top-down MS experiments from human, swine, and mouse cardiac tissues.** Species, proteoform, modified forms, theoretical most abundant mass, experimental most abundant mass, and mass error for the proteoforms identified. Modified proteoforms were identified based on highly accurate intact mass measurements and MS/MS data. Abbreviations: ventricular isoform of myosin light chain 1 (MLC-1v), methionine (Met), N,N,N-trimethylation (N,N,N-trimethyl), and S-glutathionylation (SSG).

| Species | Proteoform | Modified Forms | Theo Mass (Da) | Expt Mass (Da) | Mass Error (ppm) |
|---|---|---|---|---|---|
| *Homo sapien* | MLC-1v | Met removal, N,N,N-Trimethyl | 21841.96 | 21841.94 | 0.9 |
| *Homo sapien* | MLC-1v | Met removal, N,N,N-Trimethyl, SSG | 22147.03 | 22147.00 | 1.4 |
| *Sus scrofa* | MLC-1v | Met removal, N,N,N-Trimethyl | 21766.92 | 21766.90 | 0.9 |
| *Sus scrofa* | MLC-1v | Met removal, N,N,N-Trimethyl, SSG | 22071.99 | 22071.93 | 2.7 |
| *Mus musculus* | MLC-1v | Met removal, N,N,N-Trimethyl | 22332.31 | 22332.31 | 0.3 |
| *Mus musculus* | MLC-1v | Met removal, N,N,N-Trimethyl, SSG | 22637.37 | 22637.35 | 0.9 |

### Supplementary Figures

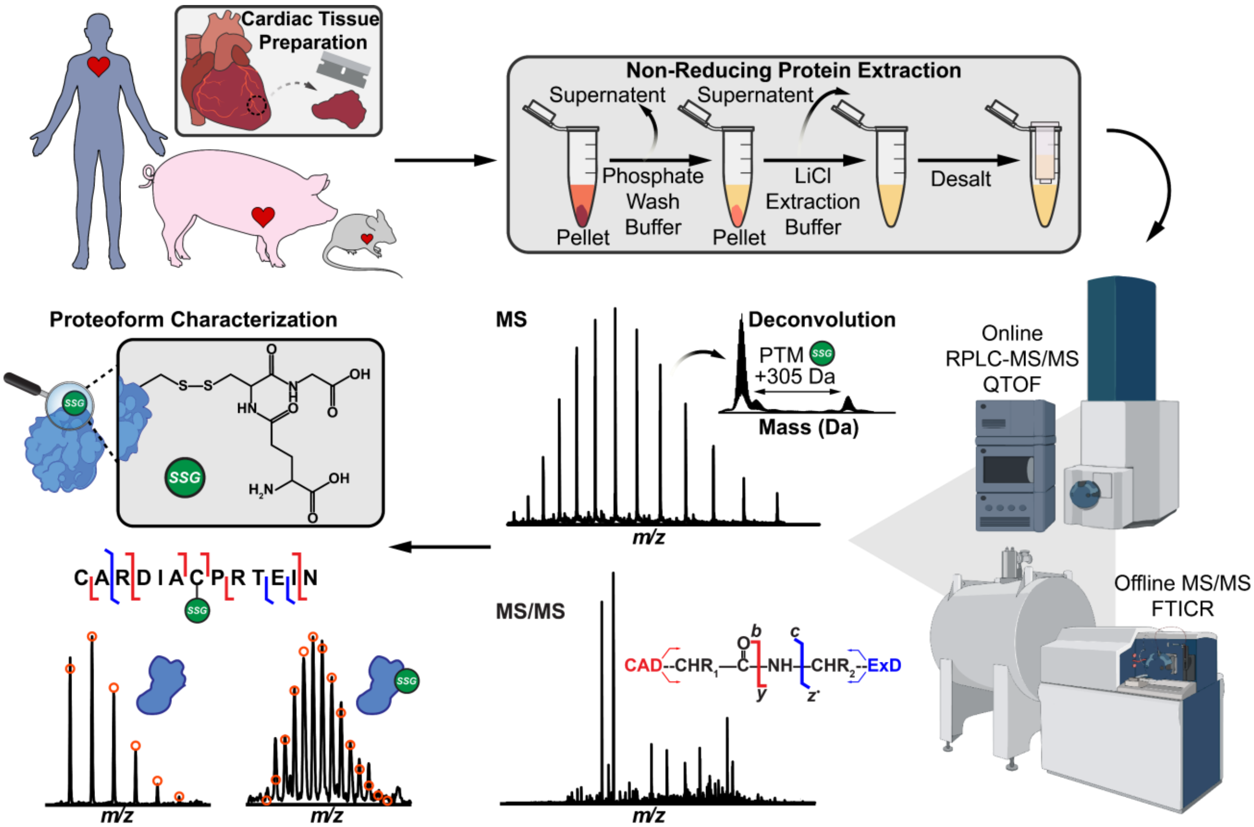


**Figure S1. Top-down MS-based proteomics workflow for the identification of protein SSG in human, swine, and mouse cardiac tissues.** Human (*n* = 12), swine (*n* = 6), and mouse (*n* = 8) cardiac tissues were first homogenized in phosphate wash buffer. Sarcomeric proteins were extracted from cardiac tissue using a non-reducing lithium chloride (LiCl) extraction buffer and then desalted using a 10 kDa molecular weight cut-off filter to prepare for mass spectrometry (MS) analysis. Intact proteins (500 ng) were separated by reverse-phase liquid chromatography (RPLC) and online MS and tandem MS (MS/MS) data were acquired using a quadrupole-time-of-flight (QTOF) mass spectrometer. Offline MS/MS data were acquired using an ultrahigh-resolution Fourier transform ion cyclotron resonance (FTICR) mass spectrometer. Protein SSG (+305.07 Da) was identified through intact mass measurements obtained from the deconvoluted mass spectra. To localize SSG, collisionally activated dissociation (CAD) and electron dissociation (ExD) fragmentation techniques were used. Proteoform characterization was performed by matching identified fragment ions to the target proteoform sequence using MASH Native v. 1.1.

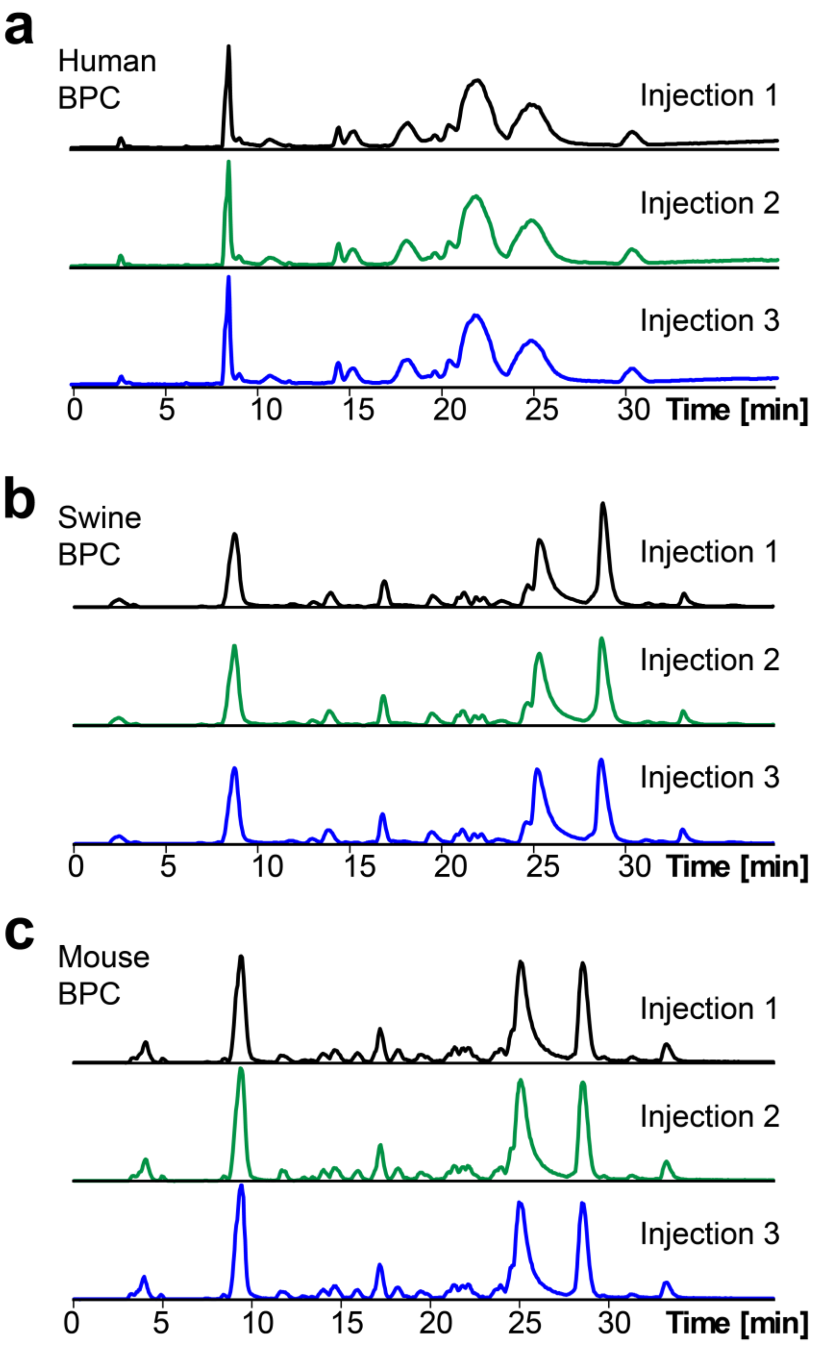


**Figure S2. Highly reproducible top-down online LC-MS/MS method.** Representative intensity normalized base peak chromatograms (BPC) for injection replicates ($n = 3$) of 500 ng of total protein from **(a)** human, **(b)** swine, and **(c)** mouse cardiac tissue lysates.

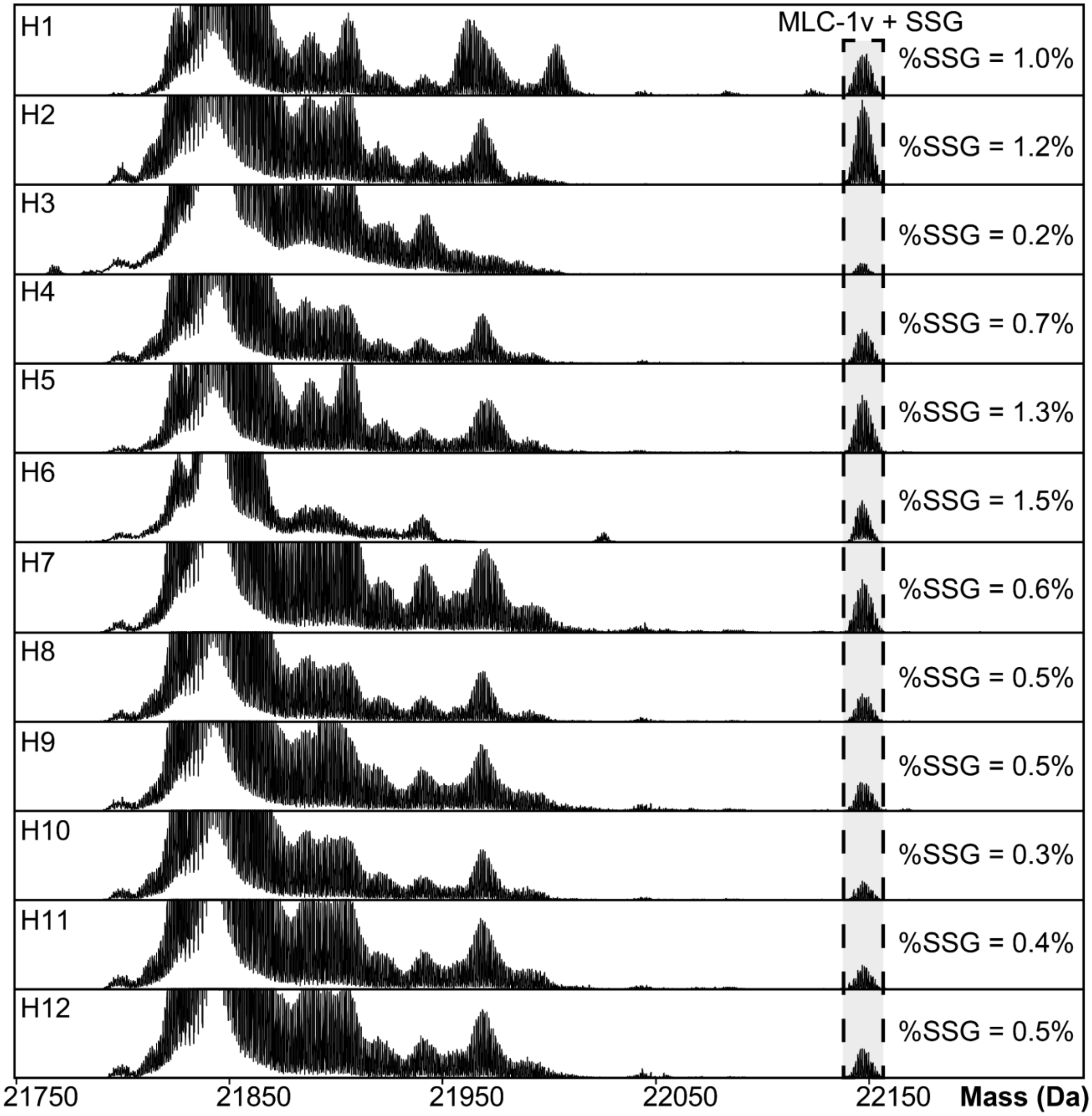


**Figure S3. Top-down proteomics identifies endogenous SSG in human MLC-1v.** Normalized deconvoluted mass spectra of MLC-1v + SSG from non-reduced human cardiac tissue lysates (*n* = 12; H1-H12) not incubated with GSSG. Total protein SSG (%SSG) was quantified based on the ratio of the deconvoluted peak intensity of the SSG proteoform to the summed peak intensities of all proteoforms of MLC-1v.

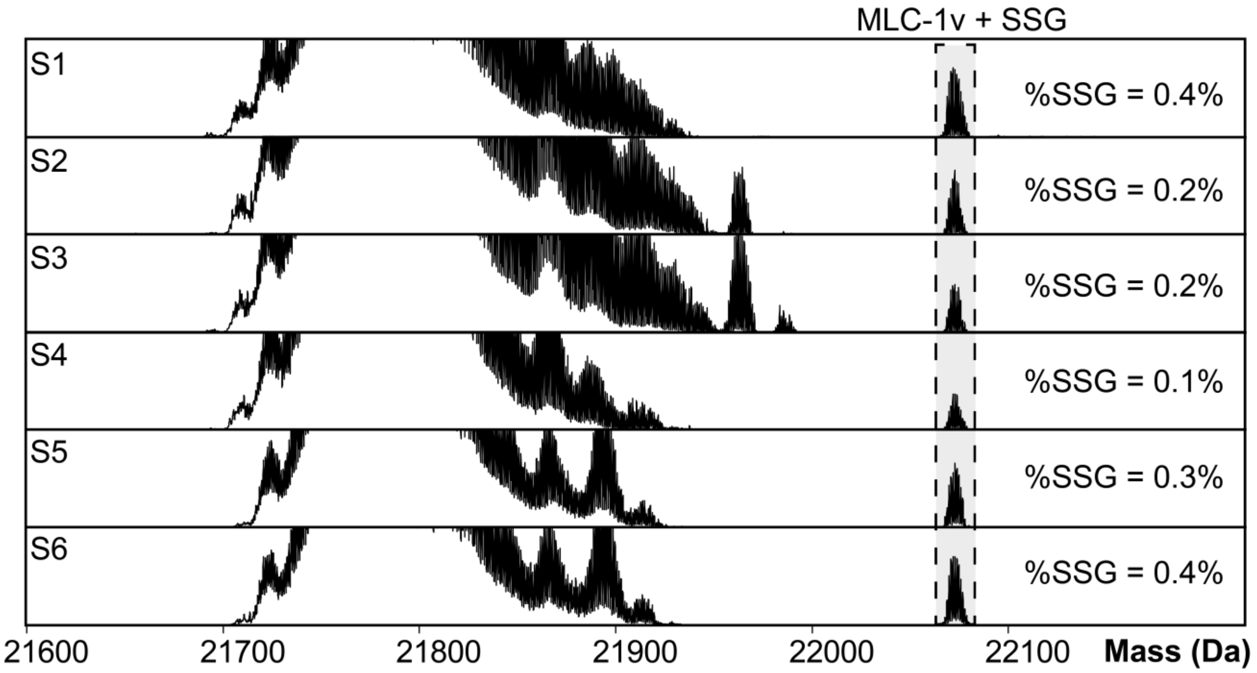


**Figure S4. Top-down proteomics identifies endogenous SSG in swine MLC-1v.** Normalized deconvoluted mass spectra of MLC-1v + SSG from non-reduced swine cardiac tissue lysates (*n* = 6; S1-S6) not incubated with GSSG. Total protein SSG (%SSG) was quantified based on the ratio of the deconvoluted peak intensity of the SSG proteoform to the summed peak intensities of all proteoforms of MLC-1v.

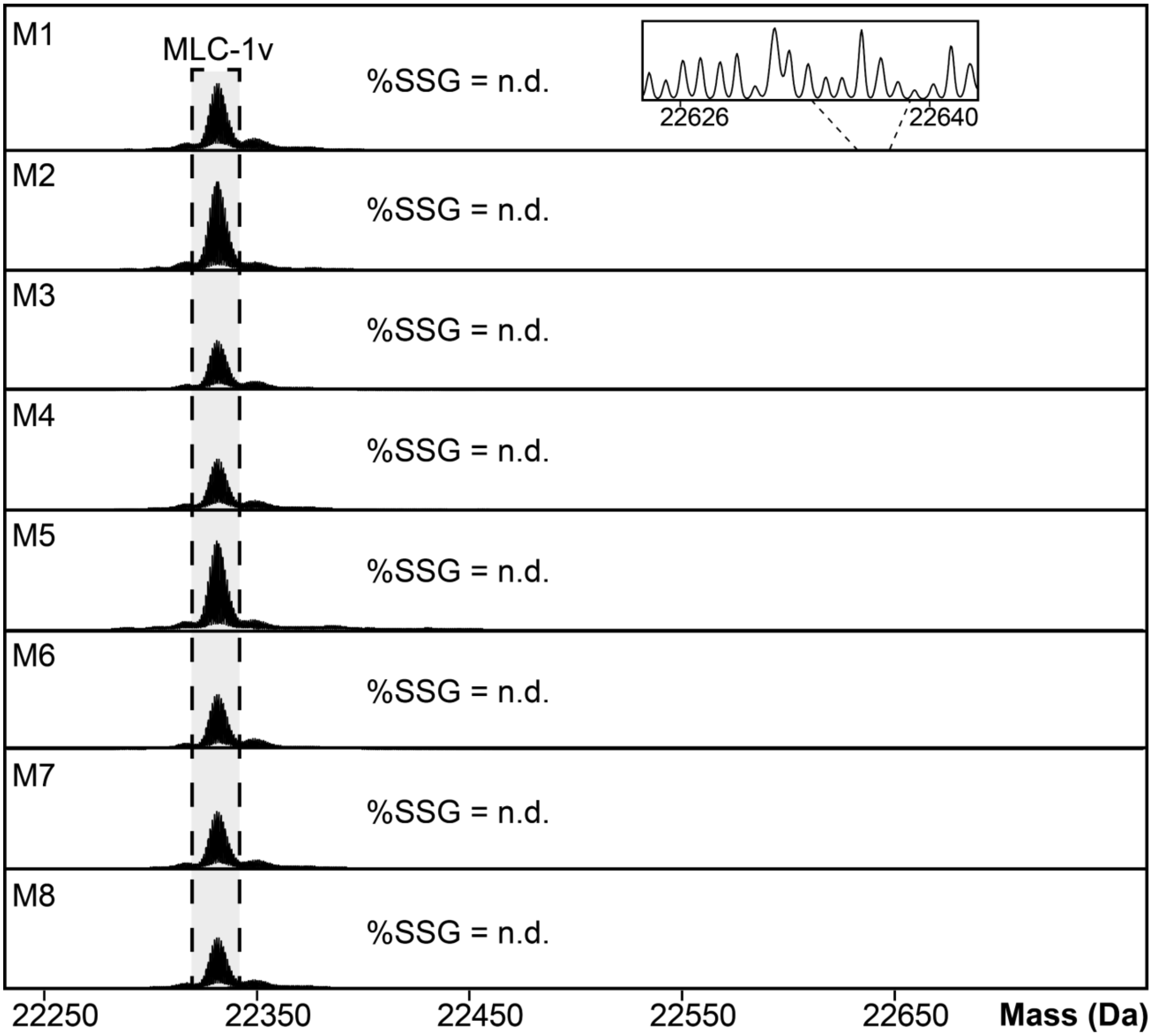


**Figure S5. Endogenous SSG is not detected by top-down proteomics in mouse MLC-1v.** Normalized deconvoluted mass spectra of MLC-1v from non-reduced mouse cardiac tissue lysates (*n* = 8; M1-M8) not incubated with GSSG. The SSG proteoform was not detected (n.d.) on MLC-1v.

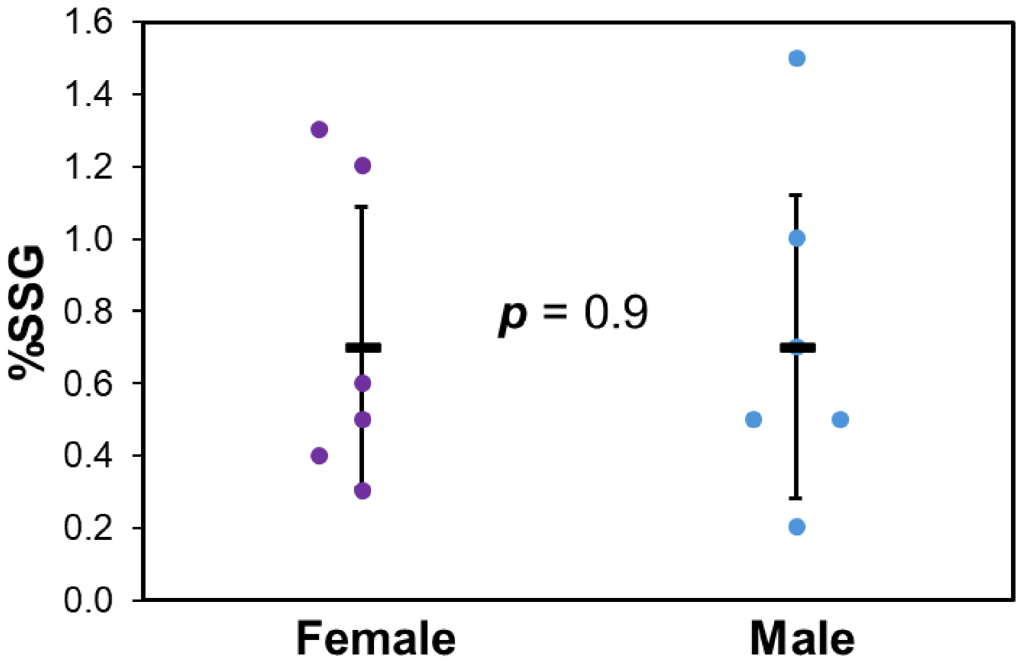


**Figure S6. Sex-based differences of SSG in human MLC-1v.** Comparison of the relative abundance of SSG (%SSG) in MLC-1v between female (purple; $n = 6$) and male (blue; $n = 6$) non-failing donor human hearts. Groups were considered significantly different by paired student t-tests with $p < 0.05$ and not significantly different with $p > 0.05$. Error bars represent the mean ± standard error of the mean (SEM).

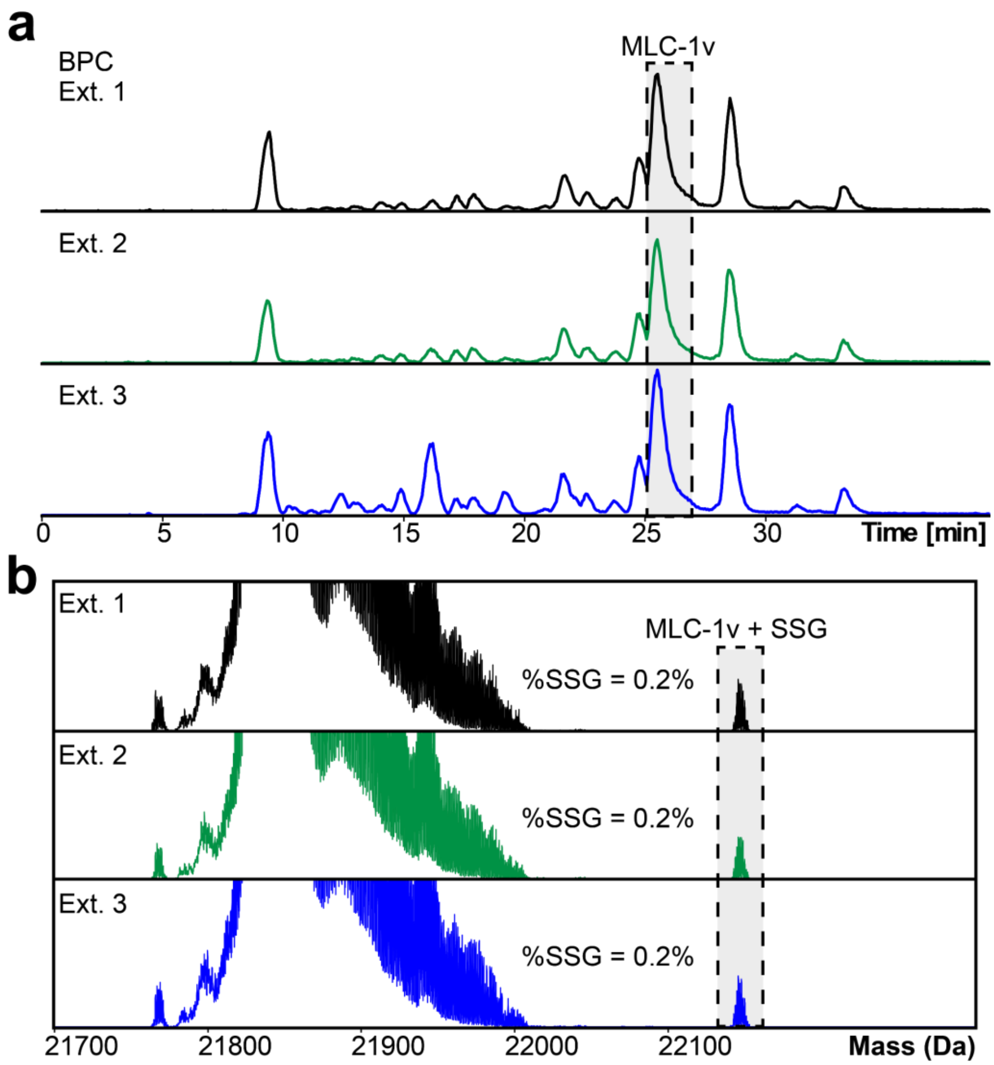


**Figure S7. Reproducible online LC-MS/MS analysis of MLC-1v from human cardiac tissue. (a)** Normalized base peak chromatograms (BPC) of individual extraction replicates (*n* = 3) from human cardiac tissue showing high reproducibility of the method. **(b)** Deconvoluted mass spectra of the MLC-1v + SSG proteoform in individual extraction replicates (*n* =3) from human cardiac tissue. Mass spectra were normalized to the same intensity. Total protein SSG (%SSG) was quantified based on the ratio of the deconvoluted peak intensity of the SSG proteoform to the summed peak intensities of all proteoforms of MLC-1v.

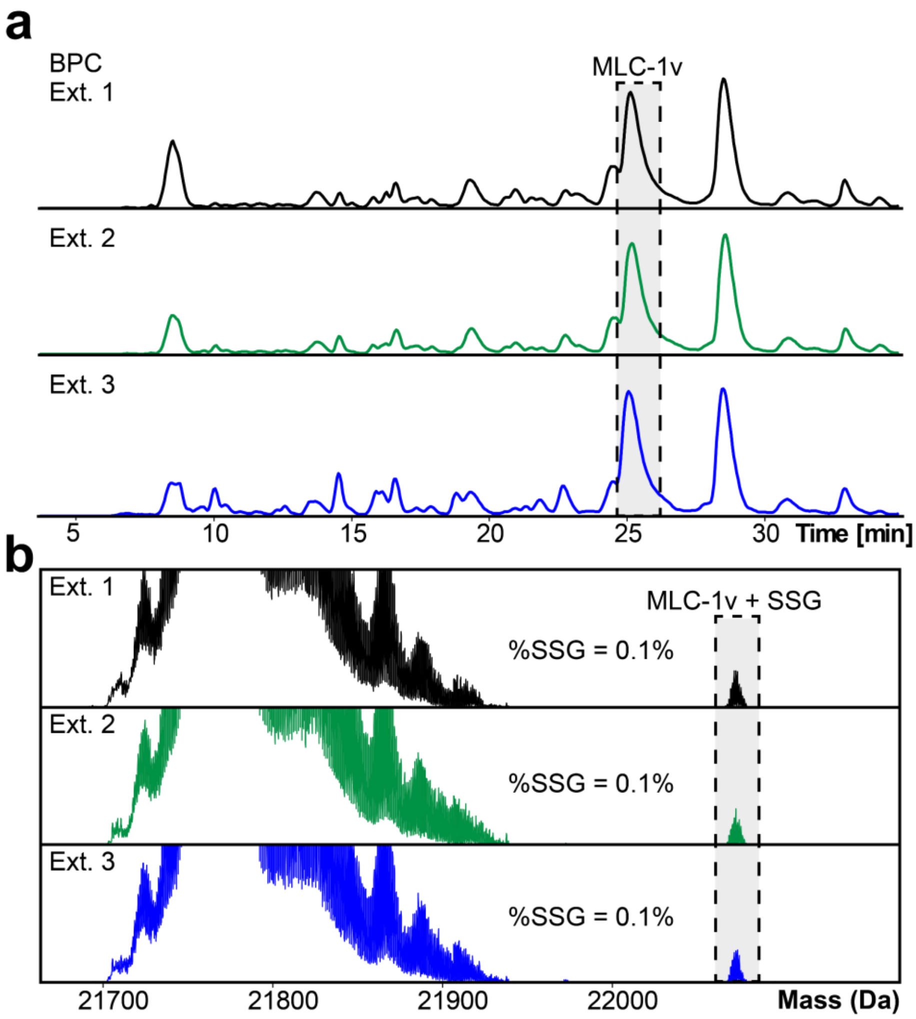


**Figure S8. Reproducible online LC-MS/MS analysis of MLC-1v from swine cardiac tissue.** **(a)** Normalized base peak chromatograms (BPC) of individual extraction replicates ($n$ = 3) from swine cardiac tissue showing high reproducibility of the method. **(b)** Deconvoluted mass spectra of the MLC-1v + SSG proteoform in individual extraction replicates ($n$ =3) from swine cardiac tissue. Mass spectra were normalized to the same intensity. Total protein SSG (%SSG) was quantified based on the ratio of the deconvoluted peak intensity of the SSG proteoform to the summed peak intensities of all proteoforms of MLC-1v.

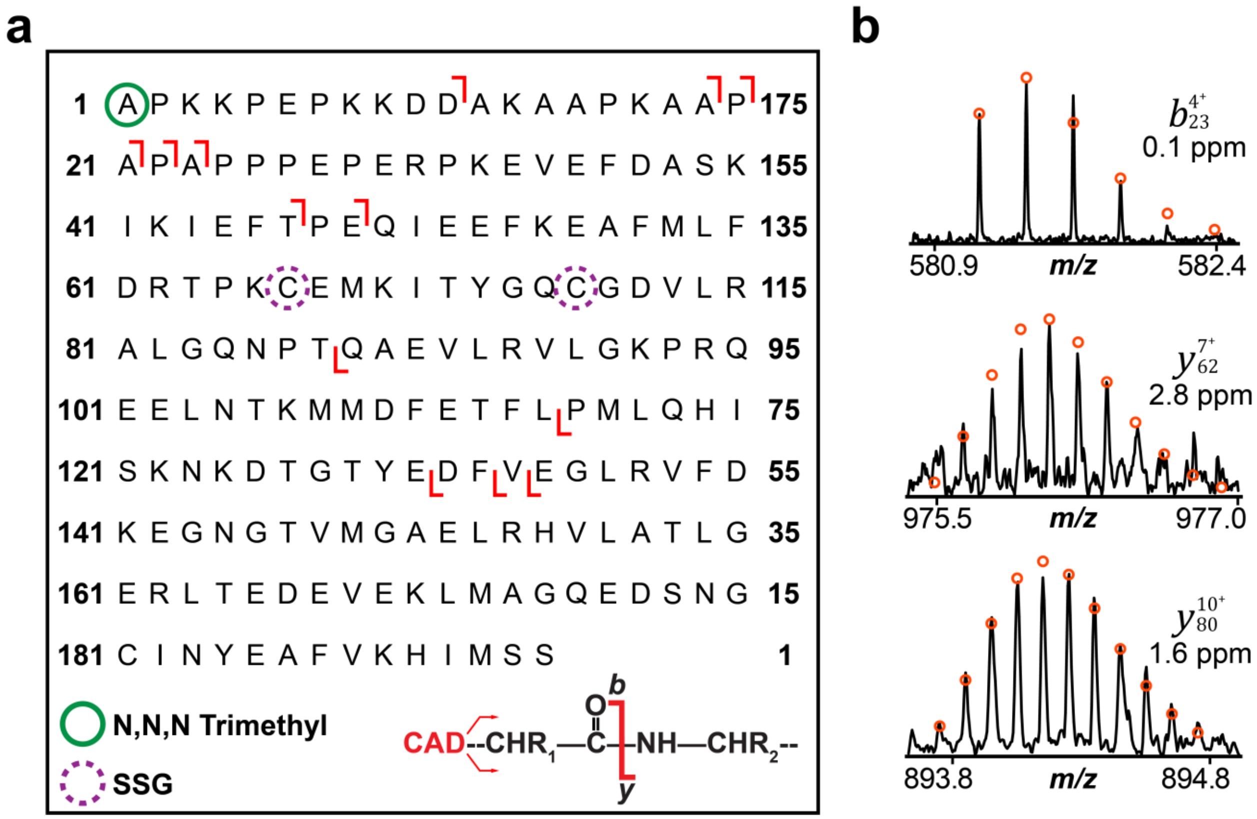


**Figure S9. Characterization of endogenous human MLC-1v + SSG using top-down proteomics.** Online LC-MS/MS analysis of the human MLC-1v + SSG proteoform (no incubation with GSSG). The MLC-1v + SSG proteoform was isolated and fragmented using collisionally activated dissociation (CAD). **(a)** The CAD fragmentation map for human MLC-1v shows SSG is confirmed to be modified on either Cys66 or Cys75. The green circle represents N,N,N-trimethylation, while the dashed purple circle represents potential SSG modification sites. **(b)** Representative CAD fragment ions. All individual ion assignments are within 10 ppm of the theoretical mass, and the theoretical isotopic distributions are indicated by the red circles. For human MLC-1v + SSG there were 8 *b* ions and 5 *y* ions achieving 7% sequence coverage.

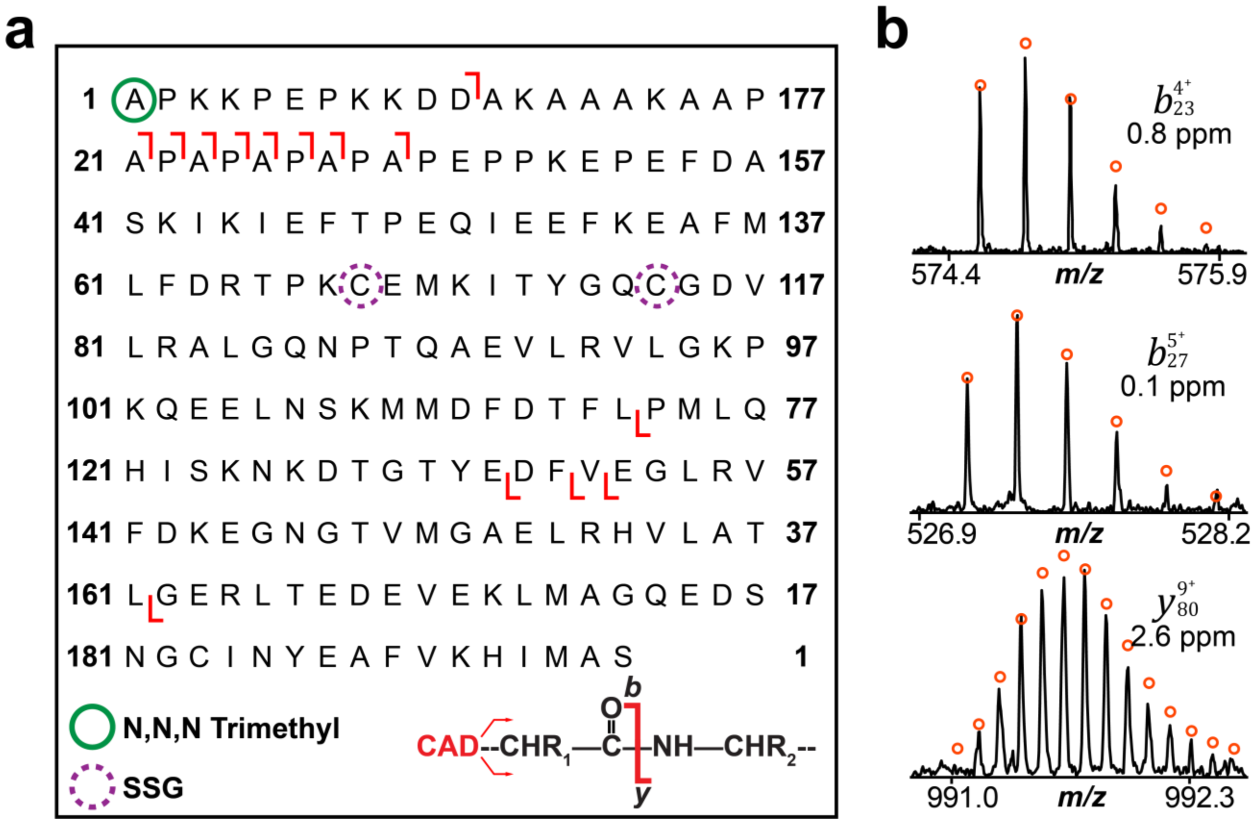


**Figure S10. Characterization of endogenous swine MLC-1v + SSG using top-down proteomics.** Online LC-MS/MS analysis of the swine MLC-1v + SSG proteoform (no incubation with GSSG). The MLC-1v + SSG proteoform was isolated and fragmented using collisionally activated dissociation (CAD). **(a)** The CAD fragmentation map for swine MLC-1v shows SSG is confirmed to be modified on either Cys68 or Cys77. The green circle represents N,N,N-trimethylation, while the dashed purple circle represents potential SSG sites. **(b)** Representative CAD fragment ions. All individual ion assignments are within 10 ppm of the theoretical mass, and the theoretical isotopic distributions are indicated by the red circles. For swine MLC-1v + SSG there were 9 *b* ions and 5 *y* ions achieving 7% sequence coverage.

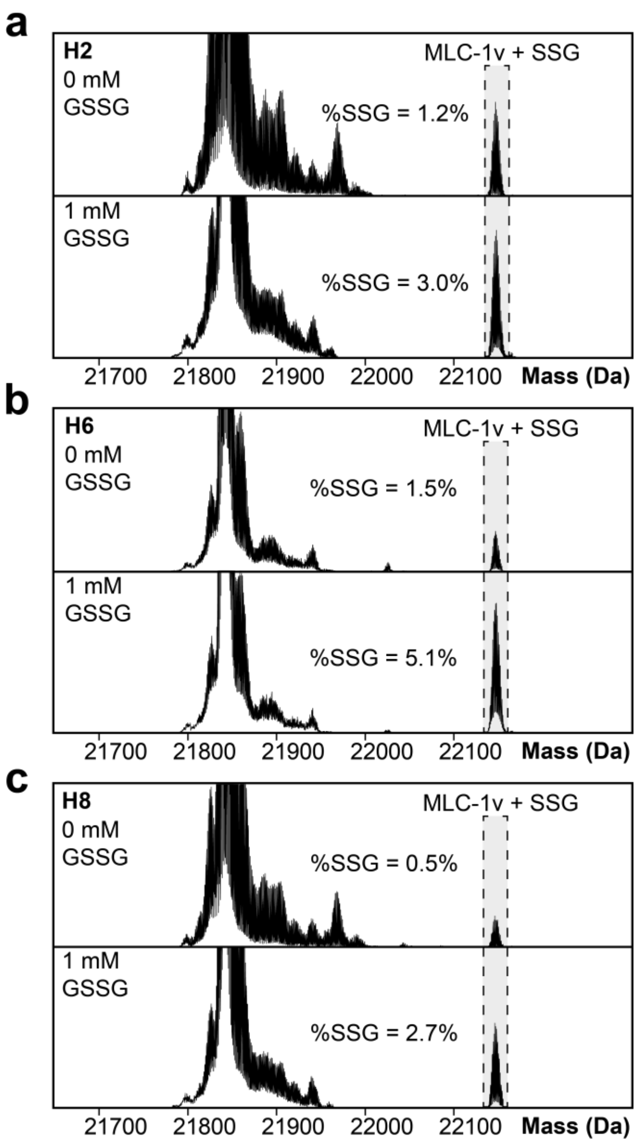


**Figure S11. Top-down proteomics reveals treatment of non-reduced human cardiac tissue lysates with GSSG increases SSG levels in MLC-1v.** Normalized deconvoluted mass spectra of the human MLC-1v + SSG proteoform in three biological replicates **(a)** H2 **(b)** H6 and **(c)** H8 that were incubated with 0 mM and 1 mM of GSSG at 4°C for 1 hour. Total protein SSG (%SSG) was quantified based on the ratio of the deconvoluted peak intensity of the SSG proteoform to the summed peak intensities of all proteoforms of MLC-1v.

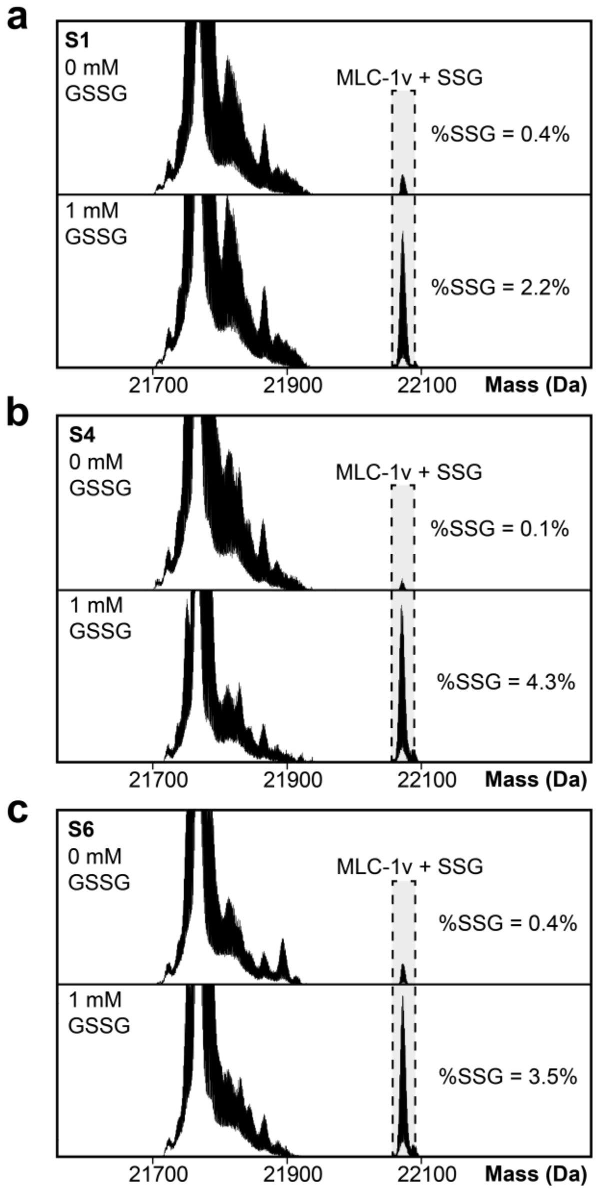


**Figure S12. Top-down proteomics reveals treatment of non-reduced swine cardiac tissue lysates with GSSG increases SSG levels in MLC-1v.** Normalized deconvoluted mass spectra of the swine MLC-1v + SSG proteoform in three biological replicates **(a)** S1 **(b)** S4 and **(c)** S6 that were incubated with 0 mM and 1 mM of GSSG at 4°C for 1 hour. Total protein SSG (%SSG) was quantified based on the ratio of the deconvoluted peak intensity of the SSG proteoform to the summed peak intensities of all proteoforms of MLC-1v.

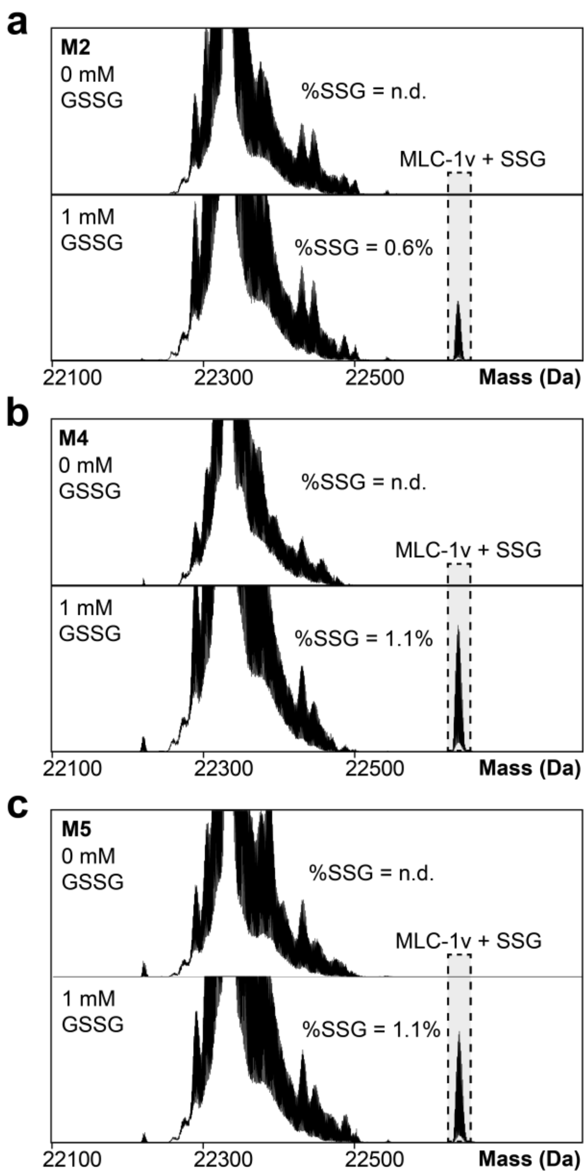


**Figure S13. Top-down proteomics reveals treatment of non-reduced mouse cardiac tissue lysates with GSSG increases SSG levels in MLC-1v.** Normalized deconvoluted mass spectra of the mouse MLC-1v + SSG proteoform in three biological replicates **(a)** M2 **(b)** M4 and **(c)** M5 that were incubated with 0 mM and 1 mM of GSSG at 4°C for 1 hour. Total protein SSG (%SSG) was quantified based on the ratio of the deconvoluted peak intensity of the SSG proteoform to the summed peak intensities of all proteoforms of MLC-1v.

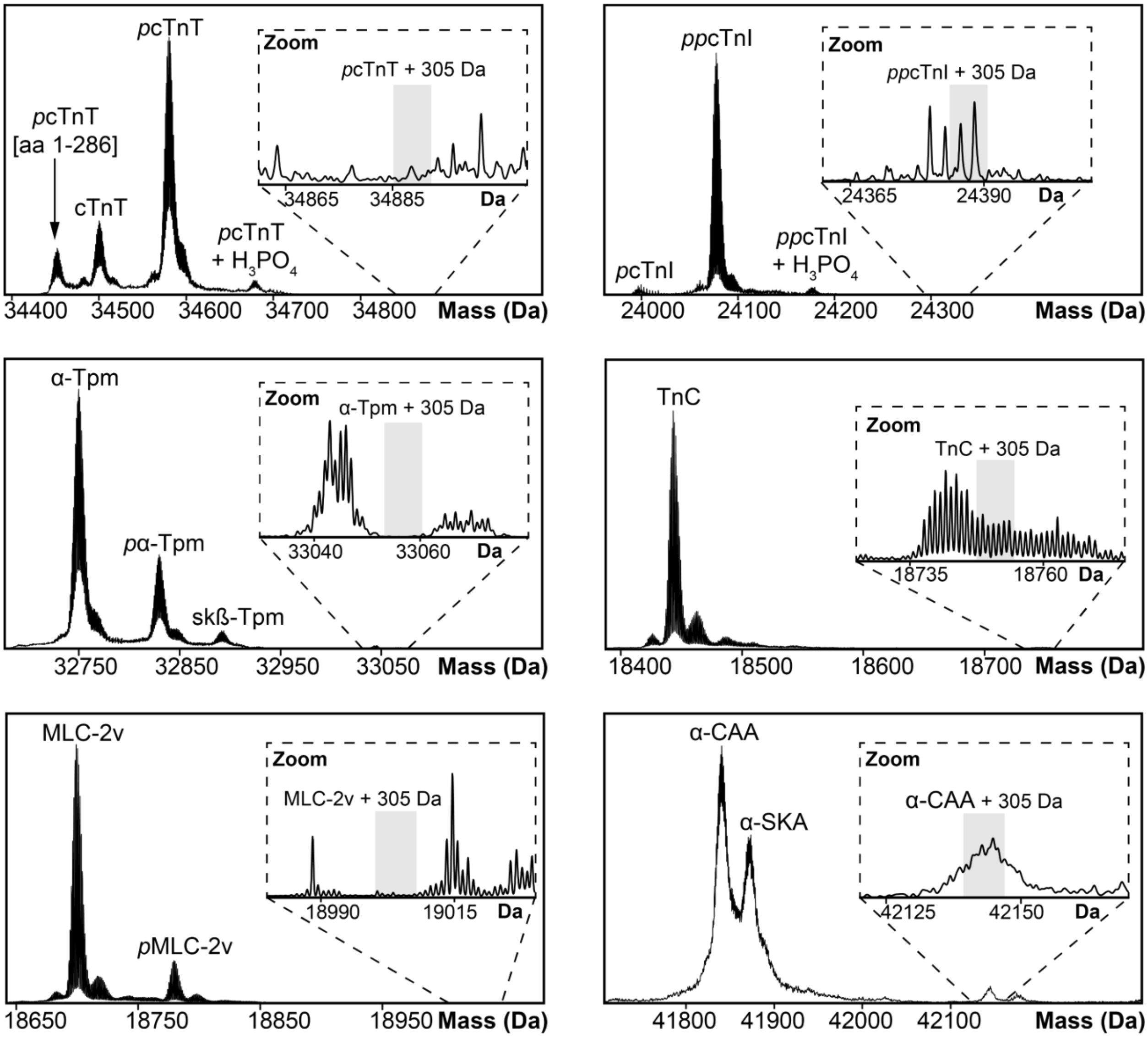


**Figure S14. Overview of major cardiac proteoforms accessed by top-down proteomics when non-reduced human tissue lysate was incubated with 1 mM GSSG.** Representative deconvoluted mass spectra showing proteoforms of major human cardiac proteins; cardiac troponin T (cTnT), cardiac troponin I (cTnI), alpha-tropomyosin (α-Tpm), troponin C (TnC), ventricular isoform of myosin light chain 2 (MLC-2v), alpha-cardiac actin (α-CAA), and alpha-skeletal actin (α-SKA). Mono- and bis-phosphorylated proteoforms are indicated with italicized "*p*" and "*pp*", respectively. Insets show a zoomed-in view of the most abundant proteoform with a +305 Da mass shift to represent the expected mass of the SSG proteoform for each protein.

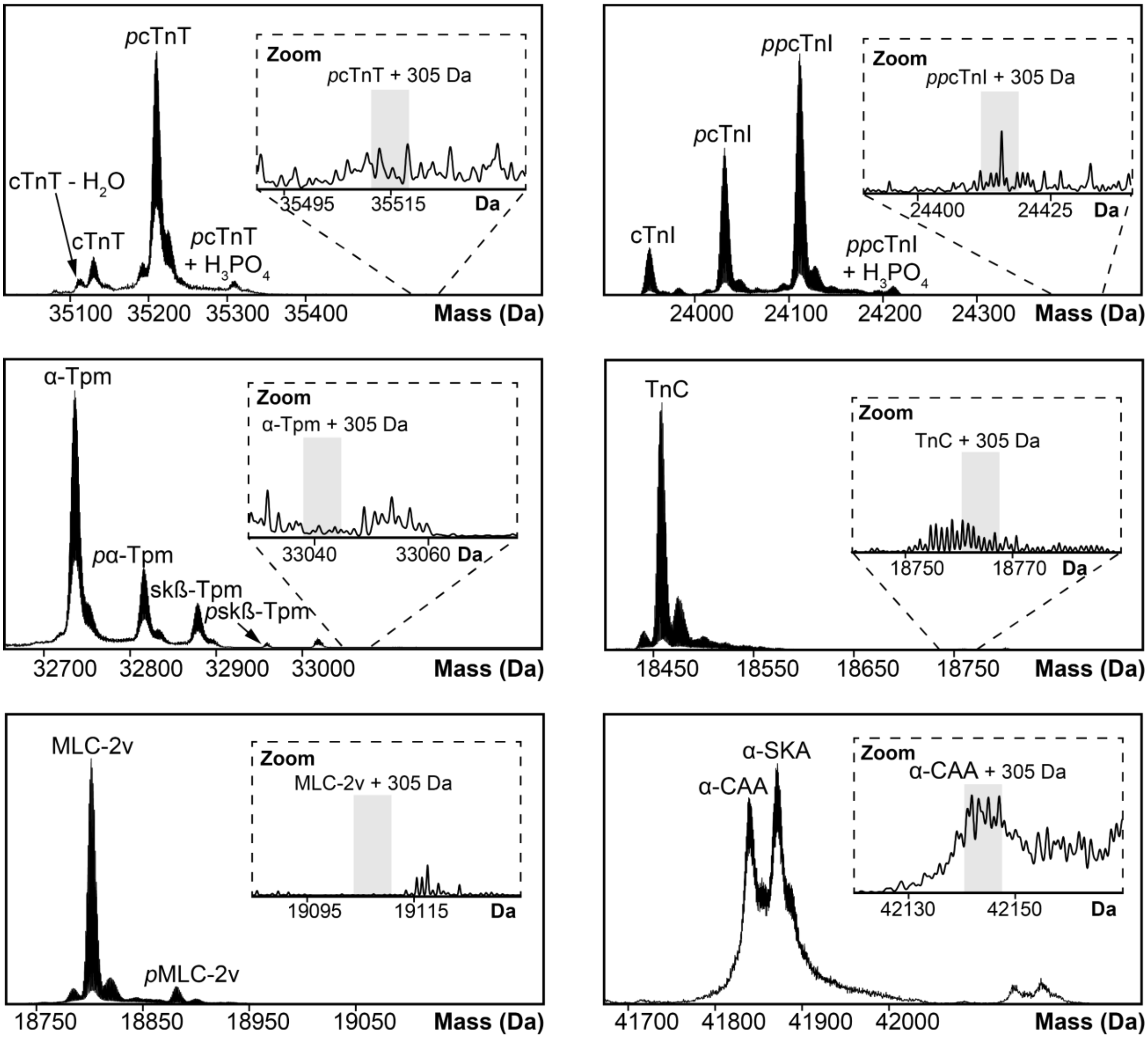


**Figure S15. Overview of major cardiac proteoforms accessed by top-down proteomics when non-reduced swine tissue lysate was incubated with 1 mM GSSG.** Representative deconvoluted mass spectra showing proteoforms of major swine cardiac proteins; cardiac troponin T (cTnT), cardiac troponin I (cTnI), alpha-tropomyosin (α-Tpm), troponin C (TnC), ventricular isoform of myosin light chain 2 (MLC-2v), alpha-cardiac actin (α-CAA), and alpha-skeletal actin (α-SKA). Mono- and bis-phosphorylated proteoforms are indicated with italicized "*p*" and "*pp*", respectively. Insets show a zoomed-in view of the most abundant proteoform with a +305 Da mass shift to represent the expected mass of the SSG proteoform for each protein.

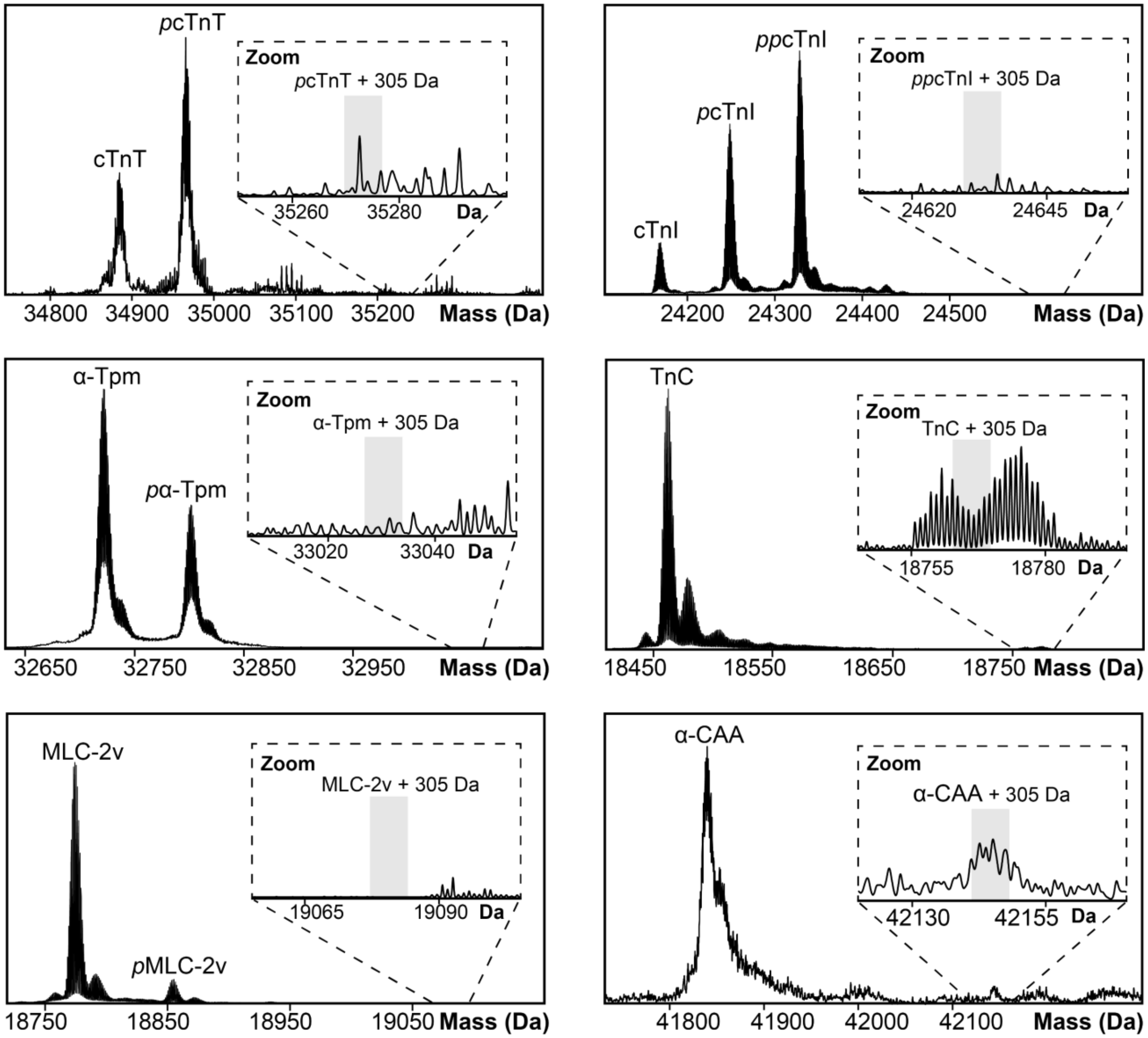


**Figure S16. Overview of major cardiac proteoforms accessed by top-down proteomics when non-reduced mouse tissue lysate was incubated with 1 mM GSSG.** Representative deconvoluted mass spectra showing proteoforms of major mouse cardiac proteins; cardiac troponin T (cTnT), cardiac troponin I (cTnI), alpha-tropomyosin (α-Tpm), troponin C (TnC), ventricular isoform of myosin light chain 2 (MLC-2v), alpha-cardiac actin (α-CAA), and alpha-skeletal actin (α-SKA). Mono- and bis-phosphorylated proteoforms are indicated with italicized "*p*" and "*pp*", respectively. Insets show a zoomed-in view of the most abundant proteoform with a +305 Da mass shift to represent the expected mass of the SSG proteoform for each protein.

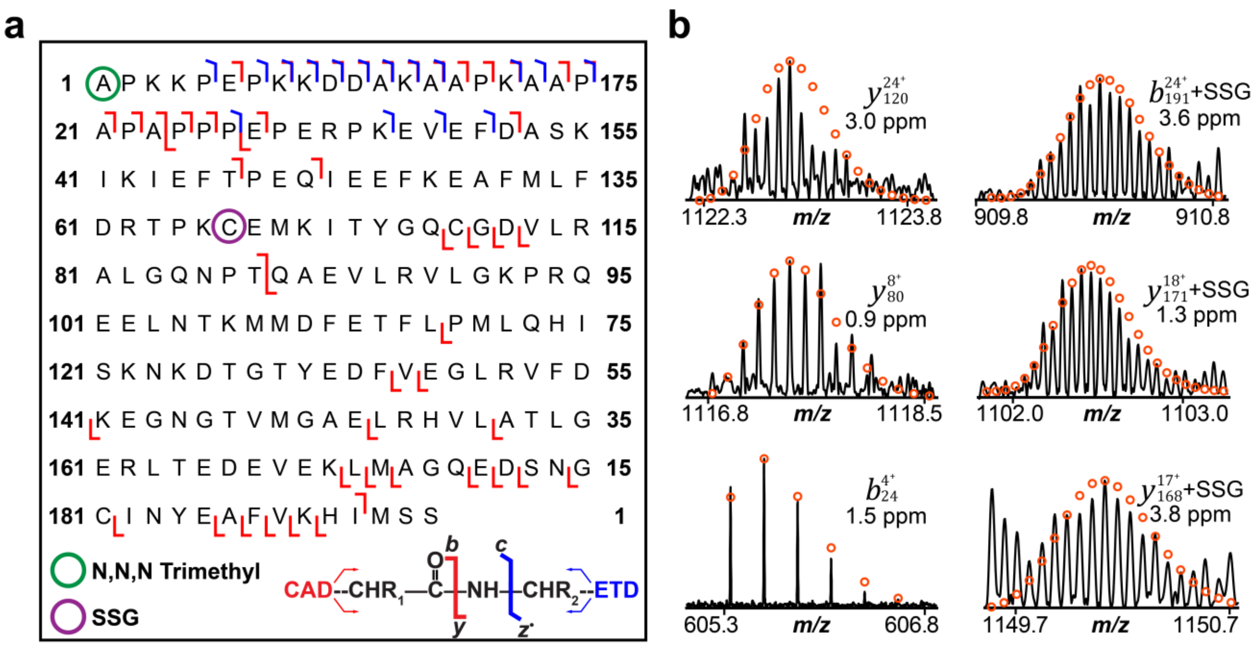


**Figure S17. Characterization of the human MLC-1v + SSG proteoform following incubation with 1 mM GSSG using top-down proteomics. (a)** Combined online and offline MS/MS fragmentation map for the MLC-1v + SSG proteoform in human cardiac tissue lysate incubated with 1 mM GSSG. MLC-1v + SSG was isolated and fragmented using collisionally activated dissociation (CAD) and electron transfer dissociation (ETD). The fragmentation map for human MLC-1v shows SSG is localized to Cys66. The green circle represents N,N,N-trimethylation, while the purple circle represents SSG. **(b)** Representative fragment ions. All individual ion assignments are within 10 ppm of the theoretical mass, and the theoretical isotopic distributions are indicated by the red circles. For human MLC-1v + SSG there were 24 *b* ions, 26 *y* ions, and 16 *c* ions achieving 29% sequence coverage.

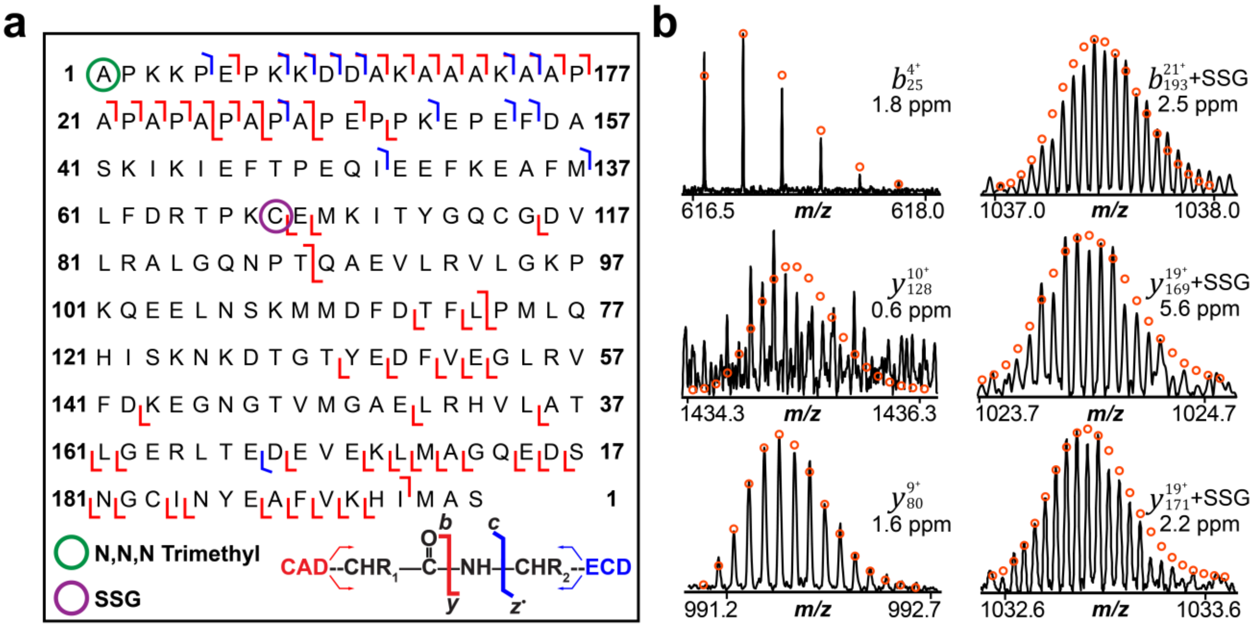


**Figure S18. Characterization of the swine MLC-1v + SSG proteoform following incubation with 1 mM GSSG using top-down proteomics. (a)** Combined online and offline MS/MS fragmentation map for the MLC-1v + SSG proteoform in swine cardiac tissue lysate incubated with 1 mM GSSG. MLC-1v + SSG was isolated and fragmented using collisionally activated dissociation (CAD) and electron capture dissociation (ECD). The fragmentation map for swine MLC-1v shows SSG is localized to Cys68. The green circle represents N,N,N-trimethylation, while the purple circle represents SSG. **(b)** Representative fragment ions. All individual ion assignments are within 10 ppm of the theoretical mass, and the theoretical isotopic distributions are indicated by the red circles. For swine MLC-1v + SSG there were 27 *b* ions, 39 *y* ions, 13 *c*, and 1 $z^{\bullet}$ ions achieving 37% sequence coverage.

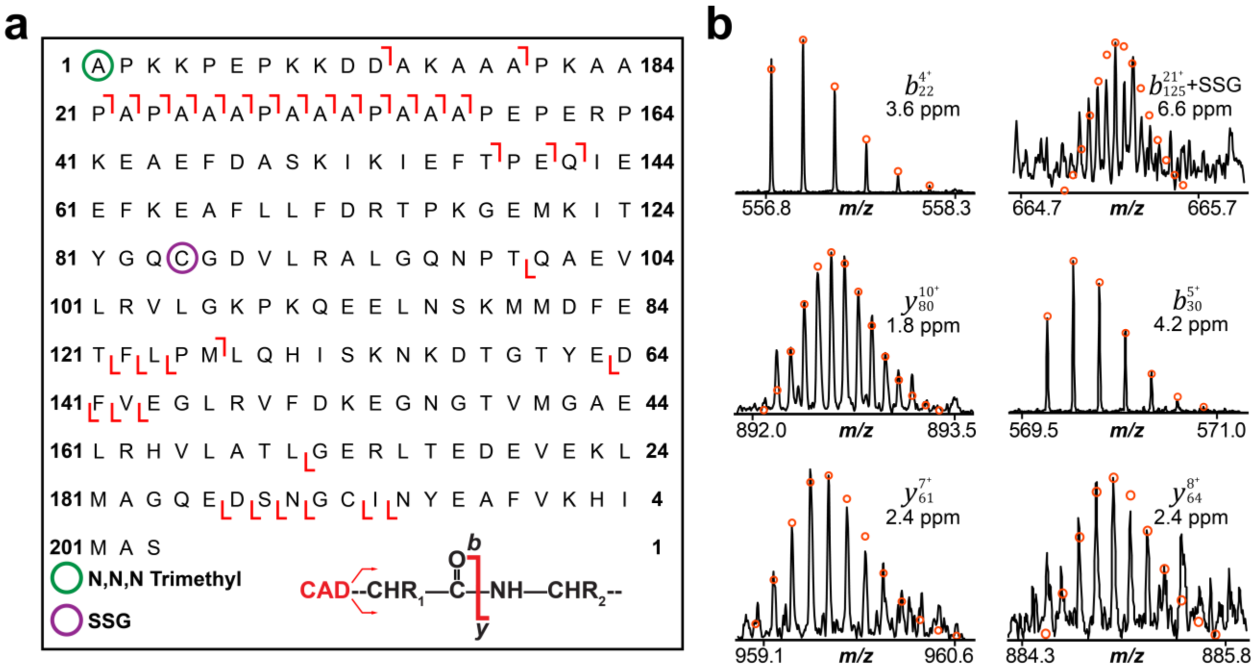

**Figure S19. Characterization of the mouse MLC-1v + SSG proteoform following incubation with 1 mM GSSG using top-down proteomics. (a)** Combined online MS/MS fragmentation map for the MLC-1v + SSG proteoform in mouse cardiac tissue lysate incubated with 1 mM GSSG. MLC-1v + SSG was isolated and fragmented using collisionally activated dissociation (CAD). The fragmentation map for mouse MLC-1v shows SSG is localized to Cys84. The green circle represents N,N,N-trimethylation, while the purple circle represents SSG. **(b)** Representative fragment ions. All individual ion assignments are within 10 ppm of the theoretical mass, and the theoretical isotopic distributions are indicated by the red circles. For mouse MLC-1v + SSG there were 20 *b* ions and 15 *y* ions achieving 17% sequence coverage.

```
sp|P08590|MYL3_HUMAN           MAPKKPEPKKDDAKAA-PKAAP--------APAPPPEPERPKEVEFDASKIKIEFTPEQI  51
sp|P09542|MYL3_MOUSE           MAPKKPEPKKDDAKAAAPKAAPAPAAAPAAAPAAAPEPERPKEAEFDASKIKIEFTPEQI  60
tr|A0A8D1HG89|A0A8D1HG89_PIG   MAPKKPEPKKDDAKAAAK-------AAPAPAPAPAPAPEPPKEPEFDASKIKIEFTPEQI  53
                               ****************              ***  * ** *** ****************

sp|P08590|MYL3_HUMAN           EEFKEAFMLFDRTPKCEMKITYGQCGDVLRALGQNPTQAEVLRVLGKPRQEELNTKMMDF  111
sp|P09542|MYL3_MOUSE           EEFKEAFLLFDRTPKGEMKITYGQCGDVLRALGQNPTQAEVLRVLGKPKQEELNSKMMDF  120
tr|A0A8D1HG89|A0A8D1HG89_PIG   EEFKEAFMLFDRTPKCEMKITYGQCGDVLRALGQNPTQAEVLRVLGKPKQEELNSKMMDF  113
                               *******:******* ********************************:*****:*****

sp|P08590|MYL3_HUMAN           ETFLPMLQHISKNKDTGTYEDFVEGLRVFDKEGNGTVMGAELRHVLATLGERLTEDEVEK  171
sp|P09542|MYL3_MOUSE           ETFLPMLQHISKNKDTGTYEDFVEGLRVFDKEGNGTVMGAELRHVLATLGERLTEDEVEK  180
tr|A0A8D1HG89|A0A8D1HG89_PIG   DTFLPMLQHISKNKDTGTYEDFVEGLRVFDKEGNGTVMGAELRHVLATLGERLTEDEVEK  173
                               :***********************************************************

sp|P08590|MYL3_HUMAN           LMAGQEDSNGCINYEAFVKHIMSS      195
sp|P09542|MYL3_MOUSE           LMAGQEDSNGCINYEAFVKHIMAS      204
tr|A0A8D1HG89|A0A8D1HG89_PIG   LMAGQEDSNGCINYEAFVKHIMAS      197
                               **********************:*
```

**Figure S20. Sequence alignment of human, swine, and mouse MLC-1v.** MLC-1v in human (UniProt ID P08590; MYL3_HUMAN), mouse (UniProt ID P09542; MYL3_MOUSE), and swine (UniProt ID A0A8D1HG89; A0A8D1HG89_PIG)**.** Human and swine MLC-1v share 95.4% sequence similarity. Human and mouse share 95.9% sequence similarity. Mouse and swine share 95.4% sequence similarity.

### Supplementary References

(1) Chapman, E. A.; Roberts, D. S.; Tiambeng, T. N.; Andrews, J.; Wang, M.-D.; Reasoner, E. A.; Melby, J. A.; Li, B. H.; Kim, D.; Alpert, A. J.; et al. Structure and dynamics of endogenous cardiac troponin complex in human heart tissue captured by native nanoproteomics. *Nature Communications* **2023**, *14* (1), 8400. DOI: 10.1038/s41467-023-43321-z.
(2) Chapman, E. A.; Li, B. H.; Krichel, B.; Chan, H.-J.; Buck, K. M.; Roberts, D. S.; Ge, Y. Native Top-Down Mass Spectrometry for Characterizing Sarcomeric Proteins Directly from Cardiac Tissue Lysate. *Journal of the American Society for Mass Spectrometry* **2024**, *35* (4), 738-745. DOI: 10.1021/jasms.3c00430.
(3) Gregorich, Z. R.; Cai, W.; Lin, Z.; Chen, A. J.; Peng, Y.; Kohmoto, T.; Ge, Y. Distinct sequences and post-translational modifications in cardiac atrial and ventricular myosin light chains revealed by top-down mass spectrometry. *Journal of Molecular and Cellular Cardiology* **2017**, *107*, 13-21. DOI: https://doi.org/10.1016/j.yjmcc.2017.04.002.
(4) Tucholski, T.; Knott, S. J.; Chen, B.; Pistono, P.; Lin, Z.; Ge, Y. A Top-Down Proteomics Platform Coupling Serial Size Exclusion Chromatography and Fourier Transform Ion Cyclotron Resonance Mass Spectrometry. *Analytical Chemistry* **2019**, *91* (6), 3835-3844. DOI: 10.1021/acs.analchem.8b04082.
(5) Peng, Y.; Gregorich, Z. R.; Valeja, S. G.; Zhang, H.; Cai, W.; Chen, Y.-C.; Guner, H.; Chen, A. J.; Schwahn, D. J.; Hacker, T. A.; et al. Top-down Proteomics Reveals Concerted Reductions in Myofilament and Z-disc Protein Phosphorylation after Acute Myocardial Infarction*. *Molecular & Cellular Proteomics* **2014**, *13* (10), 2752-2764. DOI: https://doi.org/10.1074/mcp.M114.040675.
(6) Larson, E. J.; Pergande, M. R.; Moss, M. E.; Rossler, K. J.; Wenger, R. K.; Krichel, B.; Josyer, H.; Melby, J. A.; Roberts, D. S.; Pike, K.; et al. MASH Native: A Unified Solution for Native Top-Down Proteomics Data Processing. *Bioinformatics* **2023**, btad359. DOI: 10.1093/bioinformatics/btad359.
(7) Horn, D. M.; Zubarev, R. A.; McLafferty, F. W. Automated reduction and interpretation of. *Journal of the American Society for Mass Spectrometry* **2000**, *11* (4), 320-332. DOI: 10.1016/S1044-0305(99)00157-9.
